# Human and mouse iPSC-derived astrocyte subtypes reveal vulnerability in Vanishing White Matter

**DOI:** 10.1101/523233

**Authors:** Prisca S. Leferink, Stephanie Dooves, Anne E.J. Hillen, Kyoko Watanabe, Gerbren Jacobs, Lisa Gasparotto, Paulien Cornelissen-Steijger, Marjo S. van der Knaap, Vivi M. Heine

**Affiliations:** Pediatric Neurology, Emma Children’s Hospital, Amsterdam UMC, Amsterdam, Neuroscience, Vrije Universiteit Amsterdam, The Netherlands; Department of Complex Trait Genetics, Center for Neurogenomics and Cognitive Research, Amsterdam Neuroscience, Vrije Universiteit Amsterdam, The Netherlands; Department of Functional Genomics, Center for Neurogenomics and Cognitive Research, Amsterdam Neuroscience, Vrije Universiteit Amsterdam, The Netherlands

**Keywords:** Vanishing White Matter, astrocyte subtypes, induced pluripotent stem cells, human, mouse

## Abstract

Astrocytes gained attention as important players in neurological disease, including a number of leukodystrophies. Several studies explored the generation of induced pluripotent stem cell-derived astrocytes for drug screening and regenerative studies. Developing robust models of patient induced pluripotent stem cells is challenged by high variability due to diverse genetic backgrounds and long-term culture procedures. While human models are of special interest, mouse-based models have the advantage that for them these issues are less pronounced. Here we present astrocyte differentiation protocols for both human and mouse induced pluripotent stem cells to specifically induce grey and white matter astrocytes. Both subtypes expressed astrocyte-associated markers, had typical astrocyte morphologies, and gave a reactive response to stress. Importantly, the grey and white matter-like astrocytes differed in size, complexity of processes, and expression profile, conform primary grey and white matter astrocytes. The newly presented mouse and human stem cell-based models for the leukodystrophy Vanishing White Matter replicated earlier findings, such as increased proliferation, decreased OPC maturation and modulation by hyaluronidase. We studied intrinsic astrocyte subtype vulnerability in Vanishing White Matter in both human and mouse cells. Oligodendrocyte maturation was specifically inhibited in cultures with Vanishing White Matter white matter-like astrocytes. By performing RNA sequencing, we found more differentially regulated genes in the white than in the grey matter-like astrocytes. Human and mouse astrocytes showed the same affected pathways, although human white matter-like astrocytes presented human-specific disease mechanisms involved in Vanishing White Matter. Using both human and mouse induced pluripotent stem cells, our study presents protocols to generate white and grey matter-like astrocytes, and shows astrocyte subtype-specific defects in Vanishing White Matter. While mouse induced pluripotent stem cell-based cultures may be less suitable to mimic human astrocyte subtype- or patient-specific changes, they might more robustly represent disease mutation-related cellular phenotypes as the cells are derived from inbred mice and the protocols are faster. The presented models give new tools to generate astrocyte subtypes for *in vitro* disease modeling and *in vivo* regenerative applications.

## Introduction

The importance of astrocytes in neurological disease is recognized increasingly, such as in Rett syndrome (Williams *et al*., 2014) and amyotrophic lateral sclerosis (Yamanaka and Komine, 2018), although their exact function(s) in health and disease remain unclear. Both mouse models and human induced pluripotent stem cell (hiPSC)-based cultures proved the central role of astrocytes in neuronal dysfunction (Oksanen *et al*., 2017; Rakela *et al*., 2018; Russo *et al*., 2018). Especially mouse transgenic studies advanced our knowledge on gene function and how single gene disruption affects cell development in neuronal and glial cell lineages (Leung and Jia, 2016). However, monogenic changes in mice do not always replicate the cellular or clinical phenotypes in human patients, which might be caused by phylogenetic differences between humans and rodents. The discovery of hiPSCs (Takahashi *et al*., 2007) greatly enhanced the possibility to study patient- and disease-specific cells. HiPSC-derived astrocytes successfully modeled neurodegenerative and neurodevelopmental disorders (Gupta *et al*., 2013; Chandrasekaran *et al*., 2016). However, hiPSC technology is challenged by high variability due to low standardization of *in vitro* protocols and due to induction of genetic changes during reprogramming and long term culturing (Falk *et al*., 2016). Furthermore, hiPSC-based models are labor-intensive, which often results in small sample sizes. As mouse pluripotent stem cell differentiations are more robust due to faster and more standardized protocols and the use of genetically similar inbred strains, mouse-based cultures are very useful. Thereby both human and mouse models present with advantages and disadvantages, and can therefore be complementary in studies of astrocyte functions in health and neurological disease.

Several studies indicate that astrocytes consist of functionally and morphologically heterogeneous populations of cells, that develop at different times and different locations in the central nervous system (CNS) under the influence of different intrinsic and environmental factors (Bayraktar *et al*., 2014; Molofsky and Deneen, 2015). Neurological disorders show defects in specific astrocyte subtypes (Martinian *et al*., 2009; Middeldorp *et al*., 2009; Diaz-Amarilla *et al*., 2011). This is especially clear in Vanishing White Matter (VWM), one of the more prevalent leukodystrophies, which is caused by mutations in the *EIF2B1-5* genes and for which no treatment is available (Van der Knaap, 2016). Previous studies showed that white matter astrocytes are selectively affected in VWM, while grey matter astrocytes are spared (Bugiani *et al*., 2011; Bugiani *et al*., 2013; Dooves *et al*., 2016; Leferink *et al*., 2017). These findings indicate that it is essential to generate iPSC models for disease-related astrocyte subtypes. Even though astrocyte differentiation protocols are available for both mouse (Kuegler *et al*., 2012; Kleiderman *et al*., 2016) and human stem cells (Krencik and Zhang, 2011; Emdad *et al*., 2012; Shaltouki *et al*., 2013; Tcw *et al*., 2017; Nadadhur *et al*., 2018), these do not generate specific astrocyte subtypes. Consequently, no disease modeling studies have been performed comparing different astrocyte subtypes. We are therefore in need of iPSC differentiation protocols to generate specific astrocyte subtypes, such as grey and white matter-specific astrocytes, to mimic disease-specific phenotypes.

In this study, we created *in vitro* models for VWM using hiPSCs and miPSCs (Fig. 1). We thereby present new protocols to generate iPSC-derived astrocyte subtypes, using signaling molecules stimulating grey (using fetal bovine serum; FBS) and white (incl. ciliary neurotrophic factor; CNTF) matter. The cells were characterized for astrocyte-associated morphologies, marker expression, reactivity, and were analyzed for transcriptomics using RNA sequencing (RNAseq). Interestingly, both hiPSC- and miPSC-derived white matter-like astrocytes showed specific vulnerability to VWM mutations. RNAseq analysis on hiPSC-derived astrocytes indicated differential gene expression in VWM compared to control astrocytes involved in several cellular mechanisms. Of these, genes involved in the immune system and extracellular matrix also selectively came up in miPSC-derived models, demonstrating the strength of cross-species validation in finding disease mechanisms induced by monogenic changes. Our results thereby provide new tools to generate astrocyte subtypes for the study of VWM and other astrocyte-associated diseases.

**Fig. 1.**
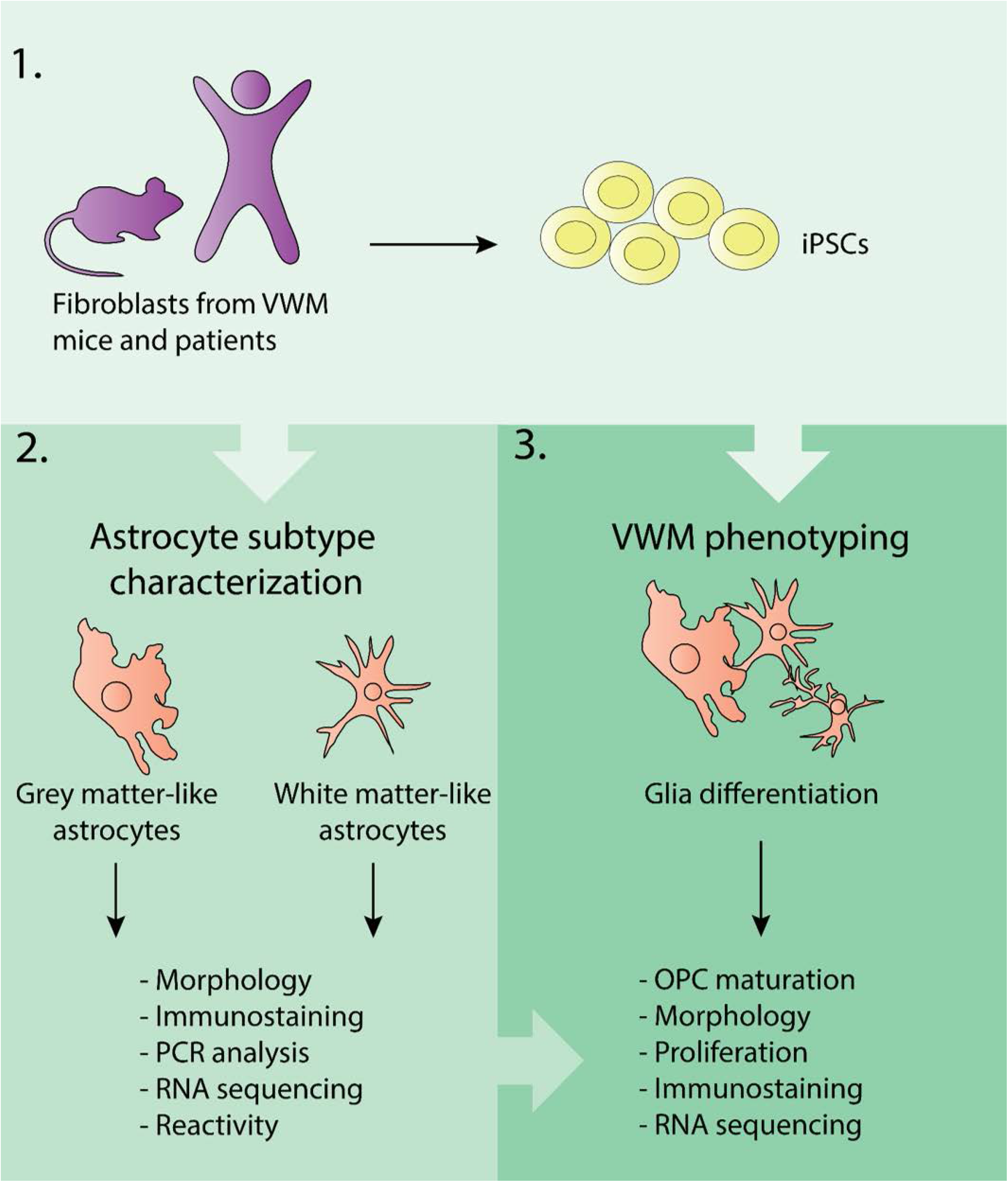
Experimental overview. Fibroblasts from human and mouse were reprogrammed to iPSCs (1) that were differentiated to grey matter-like and white matter-like astrocyte subtypes (2). To characterize these subtypes, morphology, protein and RNA expression profiles, and reactivity were assessed. The various astrocyte subtypes were also used as an *in vitro* model for VWM (3), in which OPC maturation, morphology, proliferation rate, and protein and RNA expression was studied.

## Materials and methods

### HiPSC differentiation towards astrocytes

HiPSCs were differentiated towards human astrocytes (hAstro) as described previously ((Nadadhur *et al*., 2018); see also supplementary Materials and Methods). For CNTF-hAstro subtypes, instead of using commercially available Astrocyte Medium (Sanbio, ScienCell B.V.), medium was switched to N2B27-vitA medium supplemented with EGF (20 ng/ml), FGF2 (20 ng/ml) and CNTF (20 ng/ml, Peprotech), and cells were cultured for another 2 passages in 15 days. For FBS-hAstro, the medium was switched to N2B27-vitA medium supplemented with 10% FBS, and were cultured for 2 more passages in 15 days.

### MiPSC differentiation towards mixed glial cells and astrocytes

MiPSCs obtained from wild-type (wt), *Eif2b5^Arg191His/Arg191His^ (2b5*^ho^) and *Eif2b4^Arg484Trp/Arg484Trp^Eif2b5^Arg191His/Arg191His^* (*2b4*^ho^*2b5*^ho^) mice were initially differentiated towards glial progenitor cells. The mouse models and miPSC differentiation are described in Supplementary Materials and Methods.

For mixed glial differentiation, the glial progenitor cells were subsequently cultured in mouse neural maintenance medium supplemented with 30 ng/ml T3 and 10 ng/ml NT3 from day 12 on. At day 18 cells were used for analysis.

For astrocyte differentiation cells were cultured in mouse neural maintenance medium supplemented with 10% FBS or 10 ng/ml CNTF from day 12 onwards. To obtain purer astrocyte cultures, cells were passed every 7 days for an additional 2 passages. At day 32 cells were used for analysis.

### Statistical analysis

For all experiments, results were considered significant at α = .05 after multiple test correction. RNA sequencing data was analyzed with R; all other data is analyzed with IBM SPSS statistics version 22 and Excel. Shapiro-Wilk test was used to confirm normal distribution of data and all data was tested with two-tailed tests. T-tests were used to compare FBS and CNTF astrocytes and VWM patients versus controls. To compare wild-type (wt), *2b5*^ho^ and *2b4*^ho^*2b5*^ho^ mouse data, one-way ANOVA with Dunnett post-hoc tests was used. See supplementary materials and methods for more information.

### Data availability

The data that support the findings of this study are available from the corresponding author upon reasonable request.

## Results

### MiPSC- and hiPSC-based models recapitulate OPC maturation inhibition by VWM astrocytes

Previous studies showed that astrocytes isolated from VWM mouse models impair primary mouse OPC maturation via secreted factors, suggesting that astrocytes are the primary affected cell type in VWM (Dooves *et al*., 2016). To show that iPSC-derived cell models recapitulate findings in primary cell models and to confirm that astrocytes cause cellular defects in VWM, we developed an *in vitro* miPSC model based on previous experiments with primary mouse cells. We generated miPSCs from wt, 2b5^ho^ and the severely affected *2b4*^ho^*2b5*^ho^ mice (**Fig. 2**). The miPSCs were differentiated towards glial cultures (mGlia) containing both astrocytes (~15 - 25% GFAP+ cells) (**Fig. 3A-C**) and oligodendrocytes (~30 - 40% Olig2^+^ cells) (**Fig. 3D-F, I**). Similar to previous primary mouse studies, oligodendrocyte maturation was significantly impaired in VWM mGlia, as was demonstrated by decreased numbers of MOG-expressing cells in both *2b5*^ho^ and *2b4*^ho^*2b5*^ho^ cultures (Fig. 3H; *F*(2, 5) = 7.83, *p* = .029, post-hoc Dunnett wt *vs. 2b5*^ho^ *p* = .047, wt *vs. 2b4*^ho^*2b5*^ho^ *p* = .047), and decreased numbers of MBP-expressing cells in *2b4*^ho^*2b5*^ho^ cultures (**Fig. 3 D-G**; *F*(2, 5) = 11.71 *p* = .013, post-hoc Dunnett wt *vs. 2b4*^ho^*2b5*^ho^*p* = .02), while the percentage of Olig2-positive cells remained unchanged (**Fig. 3I**). To confirm that the decreased OPC maturation was caused by astrocytes, mGlia were co-cultured with primary wt mouse astrocytes. Indeed, wt mouse astrocytes rescued the oligodendrocyte maturation defect of the mGlia (**Fig. 3J-O**), demonstrating that miPSC models recapitulate cellular models using primary cells, and confirming previous findings that VWM astrocytes are responsible for OPC maturation defects.

**Fig. 2.**
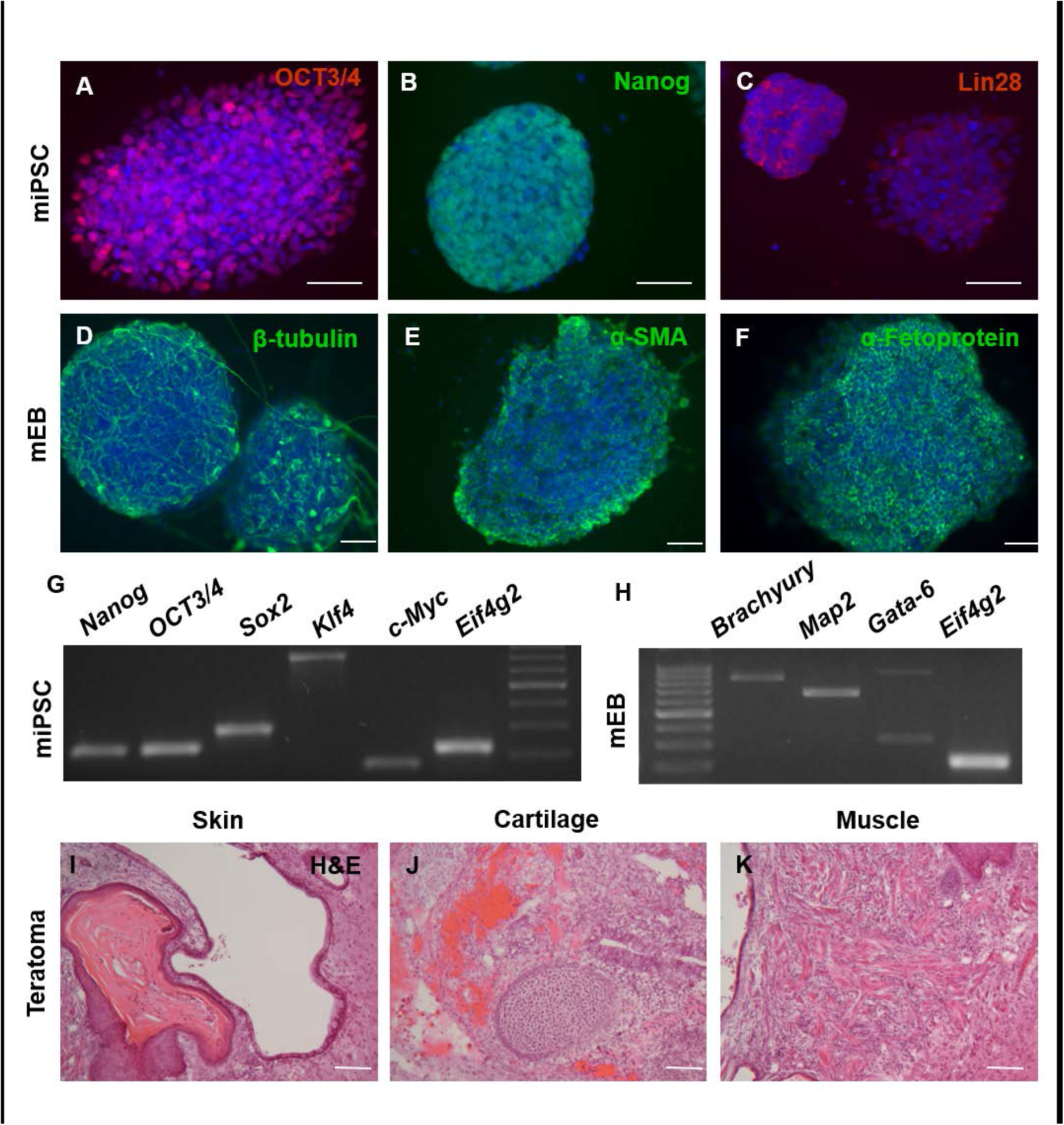
Characterization of mouse iPSC lines. Pluripotency of mouse iPSC lines was confirmed by immunostaining for pluripotency markers OCT3/4 (A), Nanog (B), and Lin28 (C), and by spontaneous differentiation into embryoid bodies after which markers of the three germ layers were present: ectoderm (β-tubulin; D), mesoderm (α-SMA; E) and endoderm (α-fetoprotein; F). RT-PCR showed mRNA expression of pluripotency markers assessed in miPSCs (G) and germ layer markers assessed in embryoid bodies (H). After injection the miPSC formed teratomas, as indicated by H&E staining for bone tissue (I), cartilage (J), and hair follicles (K). Images show an example of the characterization of one of the 2b4^ho^2b5^ho^ lines. Scalebar = 50 μm.

**Fig. 3.**
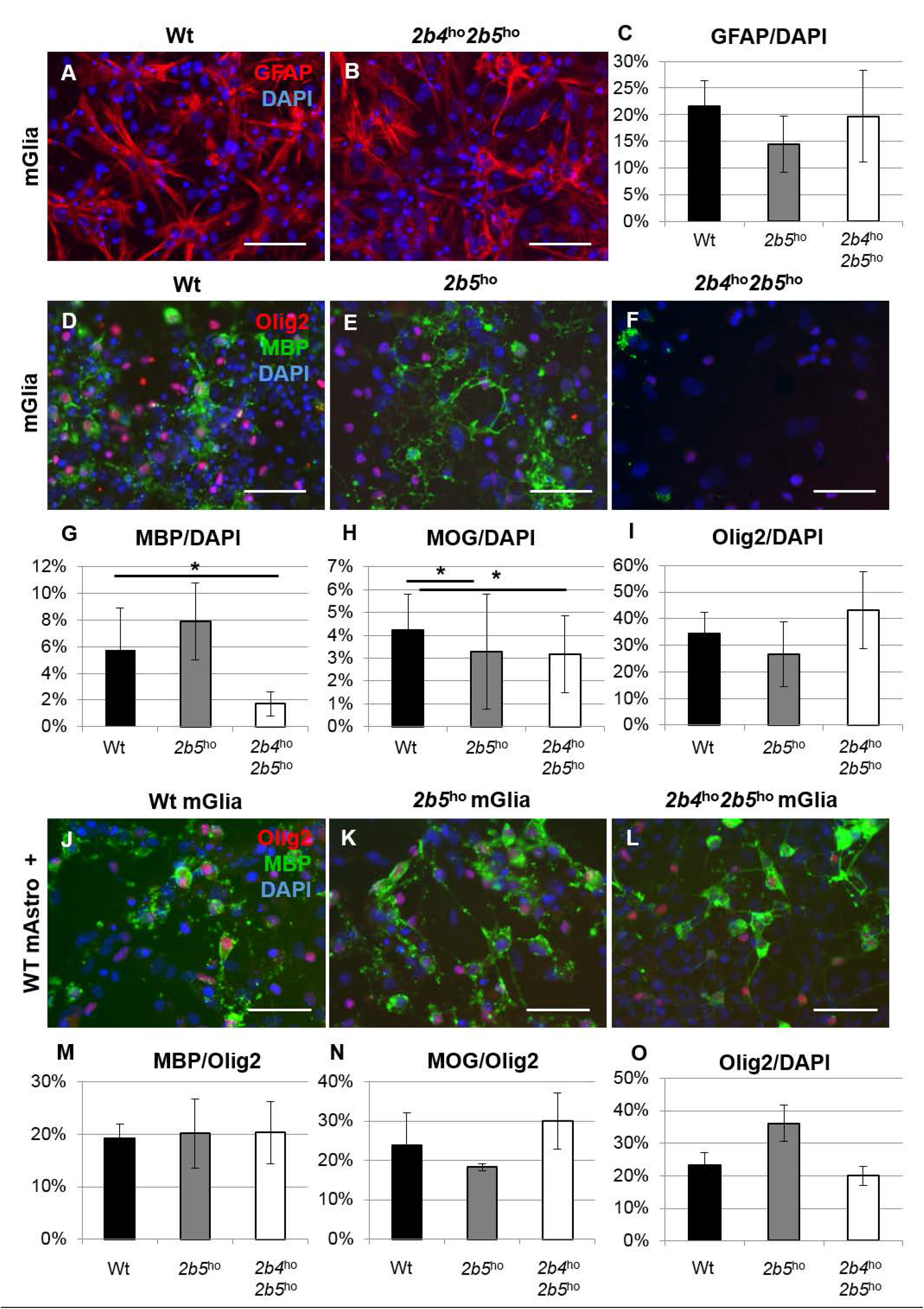
Wt astrocytes rescue the maturation defect of VWM mouse iPSC-derived oligodendrocytes. Mixed glial differentiation contained GFAP^+^ astrocytes, as shown by immunocytochemistry (A,B) and quantification (C). Oligodendrocyte maturation was addressed in wt, *2b5*^ho^ and *2b4*^ho^*2b5*^ho^ mGlia by immunostaining for MBP, MOG and Olig2 (D-I). Cell counts confirmed the decrease in the number of MBP+ (G) and MOG^+^ cells (H) in VWM without a decrease in the number of Olig2^+^ cells (I). Percentages of positive cells compared to the total number of DAPI^+^ cells are shown (wt *n* = 3; *2b5*^ho^ *n* = 3; *2b4*^ho^*2b5*^ho^ *n* = 3). To investigate the effect of healthy astrocytes on the OPC maturation defect, the wt, *2b5*^ho^, and *2b4*^ho^*2b5*^ho^ mGlia were grown on a monolayer of wt primary mouse astrocytes, and oligodendrocyte maturation was assessed using an immunostaining for MBP, MOG and Olig2 (J-O). The oligodendrocyte maturation was no longer reduced in VWM cultures, as demonstrated by no significant differences in the MBP/Olig2 (M) MOG/Olig2 (N) ratios between WT, *2b5*^ho^ and *2b4*^ho^*2b5*^ho^ mGlia. The Olig2/DAPI ratio was slightly, but not significantly, increased in 2b5^ho^ cultures (O; wt n = 3, *2b5*^ho^ n = 3; *2b4*^ho^*2b5*^ho^ n = 4). Scalebar = 50 μM. * = significant at *p* < .05. Bars in C, G-I, M-O represent mean ± SEM.

To recapitulate findings with human iPSC models, we generated hiPSCs from VWM patients (**Fig. 4**) (four iPSC lines from two patients, for extra information see supplementary M&M table 1) and healthy control donor fibroblasts (five hiPSC lines from three control donors; supplementary M&M table 1). To generate astrocytes, we differentiated control and VWM hiPSCs towards hiPSC-derived astrocytes (hAstro) as described earlier (Nadadhur *et al*., 2018). We characterized the hAstro for expression of the astrocyte-associated markers GFAP, NESTIN, Id3, CD44 and SOX9 by immunocytochemistry (**Fig. 5A-F**), and *NESTIN, BLBP, S100B, ALDOC, AQP4, GLAST, GFAP* and *CD44* by RT-PCR (**Supplementary Fig. 1A**). Functionality of hAstro was demonstrated using calcium imaging, showing glutamate uptake in both control and VWM lines (**Supplementary Fig. 1B**). To confirm increased proliferation in VWM astrocytes as was described earlier for primary human astrocytes (Bugiani *et al*., 2011), we performed a BrdU incorporation assay. The VWM hAstro showed a significant increase in the percentage of BrdU-positive cells compared to control hAstro (**Fig. 5G, H**; independent *t*(6) = 2.56, *p* = .*042*). To show that hiPSC-derived astrocytes can mediate an oligodendrocyte maturation defect, primary mouse OPCs were cultured in astrocyte-conditioned medium (ACM) from control and patient hAstro (**Fig. 5I, J**). ACM from VWM hAstro significantly impaired OPC maturation, as measured by the percentage of MBP-positive of the total numbers OLIG2-positive cells (**Fig. 5I, J, K**; independent *t*(3) = 2.34, *p* = *.049*). As hyaluronic acid (HA) has been previously described to inhibit OPC maturation (Back *et al*., 2005; Sloane *et al*., 2010) and is increased in brains of VWM patients (Bugiani *et al*., 2013), ACM from hiPSC-based cultures was treated with hyaluronidase (HYAL). Addition of HYAL significantly increased OPC maturation in VWM ACM cultures (independent *t*(3) = −8.62, *p* = *.002*), recovering towards control levels (**Fig. 5K**). Altogether, our results demonstrate that hiPSC models recapitulate findings in both primary human cells and in miPSC cell models. Furthermore, we confirmed an astrocyte-intrinsic defect in VWM using hiPSC models.

**Fig. 4.**
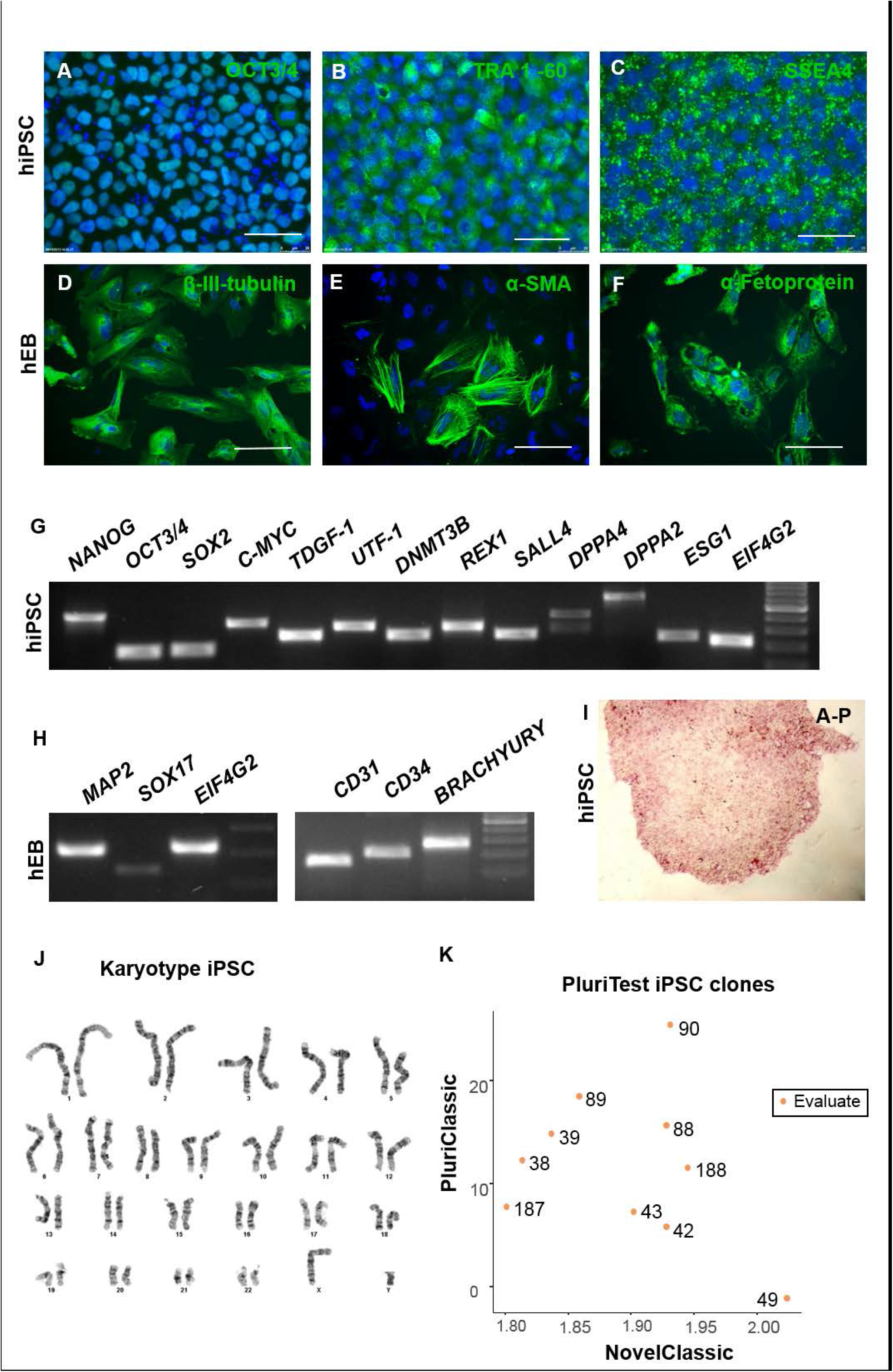
Characterization of human iPSC lines. Pluripotency of human iPSC lines is confirmed by immunostaining for pluripotency markers OCT3/4 (A), TRA-1-60 (B), and SSEA4 (C), and by spontaneous differentiation into EB resulting in expression of markers of the three germ layers ectoderm (β-tubulin; D), mesoderm (α-SMA; E) and endoderm (α-fetoprotein; F). RT-PCR analysis showed mRNA expression of pluripotency markers in hiPSCs (G), and germ layed markers after EB differentiation (H). hiPSC colonies stained positive with alkaline phosphatase (I). Karyotyping showed no chromosomal abnormalities (J), and a RNA sequencing pluritest demonstrated pluripotency of hiPSC lines (K). Images show representative examples of hiPSC lines. Scalebar = 50 μm.

**Fig. 5.**
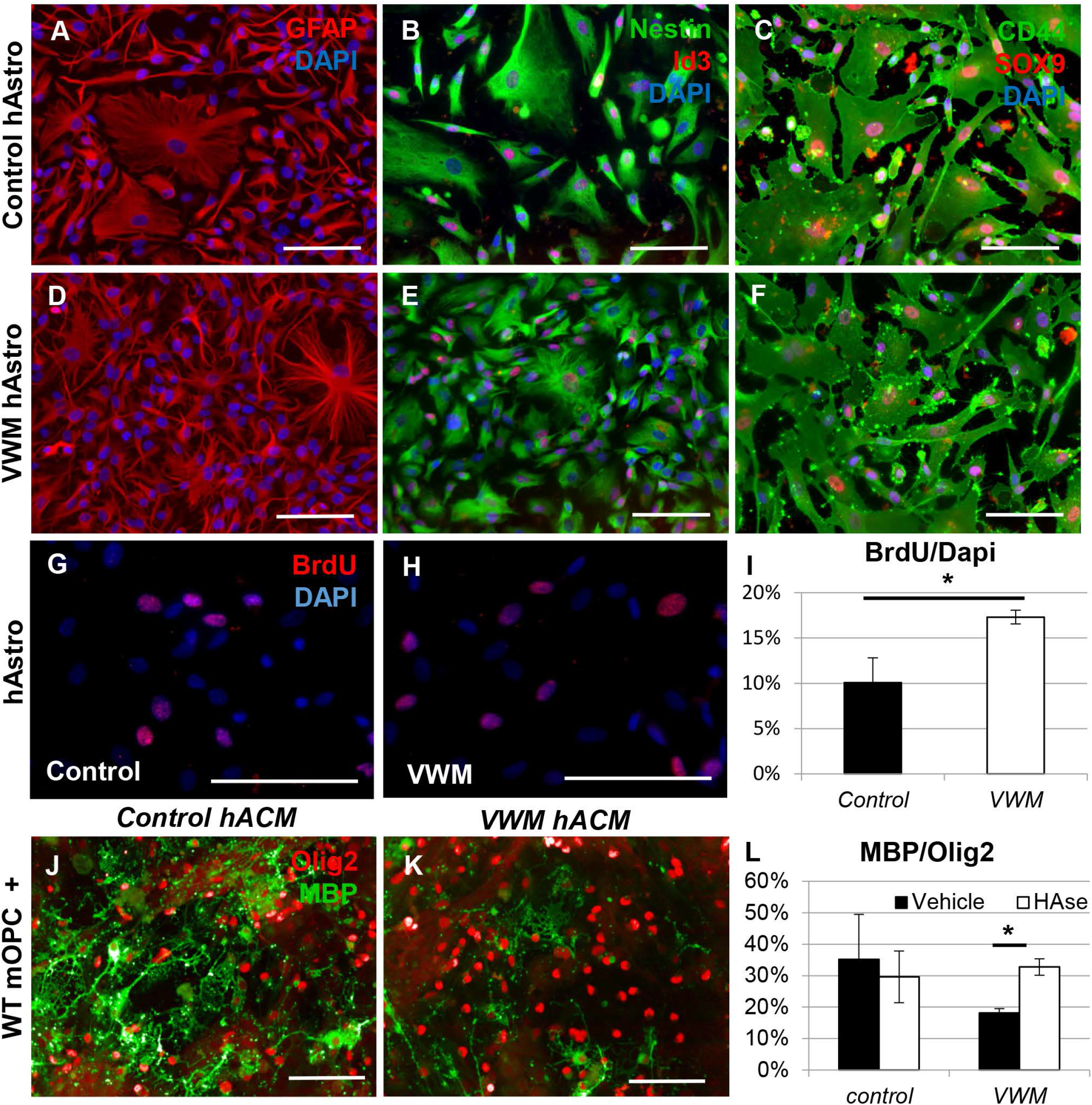
Human iPSC-derived astrocytes confirm a cell-intrinsic VWM phenotype. Immunostaining showed expression of astrocyte markers GFAP (A&D), Nestin and Id3 (B&E), CD44 and SOX9 (C&F) in both control (A–C) and VWM (D–F) hAstro. To quantify proliferation of the astrocytes, BrdU labeling was performed. A representative control (G) and VWM (H) astrocyte line is shown. Quantification of the proliferation was assessed using the percentage of BrdU-labeled cells of the total population of cells (DAPI) in Control B-E, G & I and VWM B, C, E-G, J, I hAstro lines. VWM hAstro showed increased proliferation compared to control hAstro (I). Immunostaining for Olig2 and MBP of primary wt E18 mouse OPCs cultured for 7 days in ACM of hAstro, representative examples of one VWM (H) and one control (I) line are shown. Maturation of the OPCs was assessed using a ratio of MBP+ cells of the total Olig2^+^ population, with Control A, B, E, F and VWM A, B, D, H hAstro lines used for ACM. ACM was treated with either vehicle or hyaluronidase. OPC maturation was impaired in VWM ACM as compared to control ACM, and hyaluronidase rescued OPC maturation comparable to control level (K). Scalebar = 100 μm. * = *p* <0.05; ** = *p* <0.01.

### MiPSCs differentiate into distinctive astrocytic subpopulations

An increasing number of studies confirm heterogeneity among astrocytes, such as white and grey matter-astrocytes, for which no specific iPSC-based differentiation protocols are available. CNTF-reactive astrocytes are predominantly found in the white matter (Dallner *et al*., 2002), and FBS administration is standardly used to maintain cortical astrocytes (Goursaud *et al*., 2009) leading to a flat morphology that is common for grey matter astrocytes in culture. To generate white matter- or grey matter-like astrocytes, miPSCs were first differentiated to glial precursor cells that were subsequently cultured in media supplemented with CNTF or FBS respectively. The miPSC-derived astrocyte populations showed distinct morphological properties: where CNTF-induced astrocytes (CNTF-mAstro) formed a culture of cells with many thin protrusions (**Fig. 6A**), FBS-induced astrocytes (FBS-mAstro) formed a pure population of flat and round cells with less protrusions (**Fig. 6B**). These morphological differences were quantified using automated Columbus^®^ software. The perimeter corrected for total surface area of the cell was larger in CNTF-mAstro (**Fig. 6C**, *t*(12) = 3.52, *p* = .004), consistent with morphological observations of more and thinner protrusions. FBS-mAstro had a significantly larger surface area (**Fig. 6D**, *t*(17) = −5.01, *p* < .001). While both cultures were highly GFAP-positive, the FBS-mAstro contained significantly more GFAP-positive cells (**Fig. 6E**, 97% compared to 91% in CNTF-mAstro cultures; *t*(13) = −4.29, *p* < .*001*) and Nestin-positive cells (**Fig. 6F**, 69% compared to 31% in CNTF-mAstro cultures; *t*(19) = −4.68; *p* < .*001*), showing a higher purity of FBS-induced cultures. The lower purity of CNTF-mAstro cultures was confirmed by higher levels of *Olig2* mRNA (**Fig. 6G**, *t*(12) = 2.59, *p* =.024). However, most astrocytic markers were higher expressed in CNTF-mAstro cultures on mRNA level, with significantly increased expression of *Glast* (**Fig. 6G**, *t*(14) = 5.39, *p* < .0001), *S100β* (**Fig. 6G**, *t*(16) = 2.30, *p* = .*035*), and *Gfap* (**Fig. 6G**, *t*(17) = 2.52, p = .*022*). This is consistent with findings that show that white matter astrocytes have a higher expression level of these markers (Goursaud *et al*., 2009; Hassel *et al*., 2003; Steiner *et al*., 2007). These results suggest that while both mAstro subtypes express high levels of astrocyte-associated markers and present with typical astrocyte-like morphologies, CNTF-mAstro are smaller and show increased expression of astrocyte-associated markers, in line with white matter astrocytes, whereas FBS-mAstro morphologically present as larger and rounder cells, in line with grey matter astrocytes (Lundgaard *et al*., 2014).

**Fig. 6.**
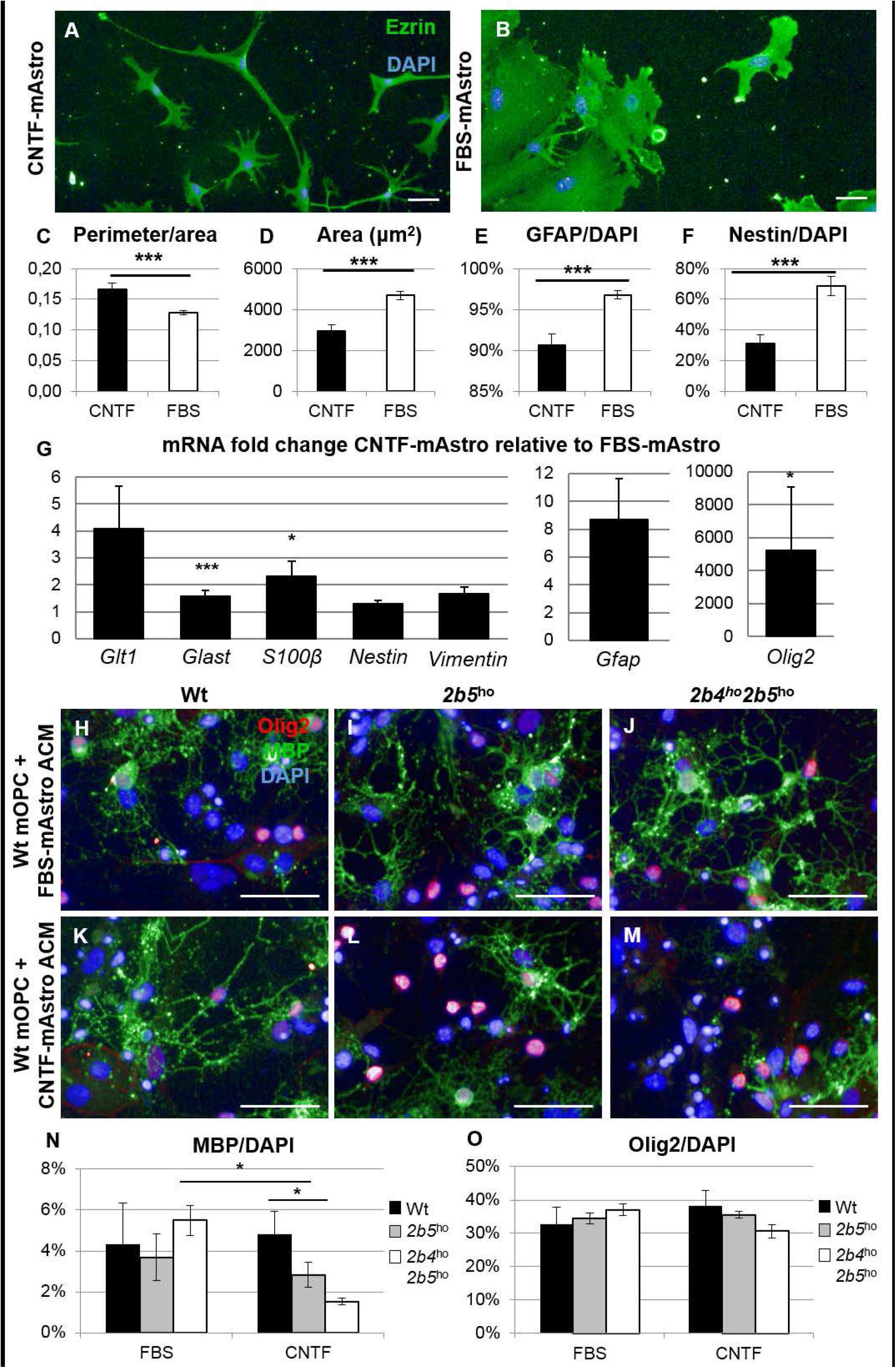
CNTF-mAstro and FBS-mAstro differ in morphology and mRNA expression level and are differentially affected by VWM mutation. Immunostaining for cytoplasmic cell surface marker Ezrin of CNTF-mAstro (A) and FBS-mAstro (B) showed morphological differences between the cells. Morphological analysis showed that CNTF-mAstro (*n* = 11) had a higher ratio of the perimeter corrected for the area (C) and had a smaller surface area (D) compared to FBS-mAstro (*n* = 10). Cell counts indicated that both populations contain > 90% of astrocytes, although the amount of GFAP+ cells was significantly lower in CNTF-mAstro compared to FBS-mAstro (E). The percentage of Nestin-expressing cells was significantly decreased in CNTF-mAstro compared to FBS-mAstro (F). mRNA expression of *Glt1, Glast, S100β, Nestin, Vimentin, Gfap* and *Olig2* was quantified in CNTF-mAstro and FBS-mAstro and is shown as mRNA fold change of CNTF-mAstro relative to FBS-mAstro (G). To assess functional defects of VWM CNTF-mAstro and FBS-mAstro, wt mouse primary OPCs were cultured in ACM collected from wt (*n* = 3), *2b5*^ho^ (*n* = 4), and *2b4*^ho^*2b5*^ho^ (*n* = 4) miPSC-derived CNTF-mAstro or FBS-mAstro (H-M). OPC maturation was quantified as the ratio of MBP^+^ oligodendrocytes to the total amount of (DAPI^+^) cells. This ratio was significantly decreased in *2b4*^ho^*2b5*^ho^ CNTF-mAstro ACM compared to wt CNTF-mAstro ACM and *2b4*^ho^*2b5*^ho^ FBS-mAstro ACM (N), while the percentage of Olig2^+^ cells was unchanged between the conditions (O). * = *p* < .05; ** = *p* < .01. Scalebar = 50 μm. Bars in C-G and N-O represent mean ± SEM.

### MiPSC-based models show selective involvement of astrocytic subtypes in VWM

To assess functional defects of VWM CNTF-mAstro and FBS-mAstro, wt mouse primary OPCs were cultured in ACM collected from wt, *2b5*^ho^, and *2b4*^ho^*2b5*^ho^ miPSC-derived CNTF-mAstro or FBS-mAstro (**Fig. 6H-M**). While ACM from VWM FBS mAstro lines did not affect OPC maturation, the ACM from *2b4*^ho^*2b5*^ho^ CNTF-mAstro significantly decreased the percentage of MBP-expressing cells compared to ACM from wt CNTF-mAstro (**Fig. 6N**; *F*(2, 8) = 6.22, *p* = .023, post-hoc Dunnett wt vs *2b4*^ho^*2b5*^ho^ *p* = .007) and compared to ACM from *2b4*^ho^*2b5*^ho^ FBS-mAstro (**Fig. 6N**, *t*(6) = −5.657, *p* = .001). The percentage of Olig2-expressing cells was unchanged between conditions (**Fig. 6O**). Altogether, these findings demonstrate that CNTF-mAstro show a higher vulnerability to VWM mutations.

Since CNTF-mAstro are selectively affected by VWM mutations, we performed transcriptome analysis on wt and *2b4*^ho^*2b5*^ho^ CNTF-mAstro to identify differentially expressed genes (DEGs) in the affected astrocytes. In total 13 genes were significantly differentially expressed between the wt and *2b4*^ho^*2b5*^ho^ CNTF-mAstro (**Supplementary Table 1**), as labelled in **Fig. 7A**. Based on enrichment analysis for DEGs with GO terms (**Supplementary Table 1**) and literature (Han *et al*., 2012; Dashzeveg *et al*., 2014; Das *et al*., 2015; Ishimoto *et al*., 2017), DEGs were overrepresented in the categories ‘Immune system’, ‘Development and proliferation’, ‘Extracellular matrix’, ‘RNA polymerase II transcription’ and ‘P53 mediated signaling’ (**Fig. 7B**), suggesting these processes are differentially regulated between VWM and wt CNTF-mAstro.

**Fig. 7.**
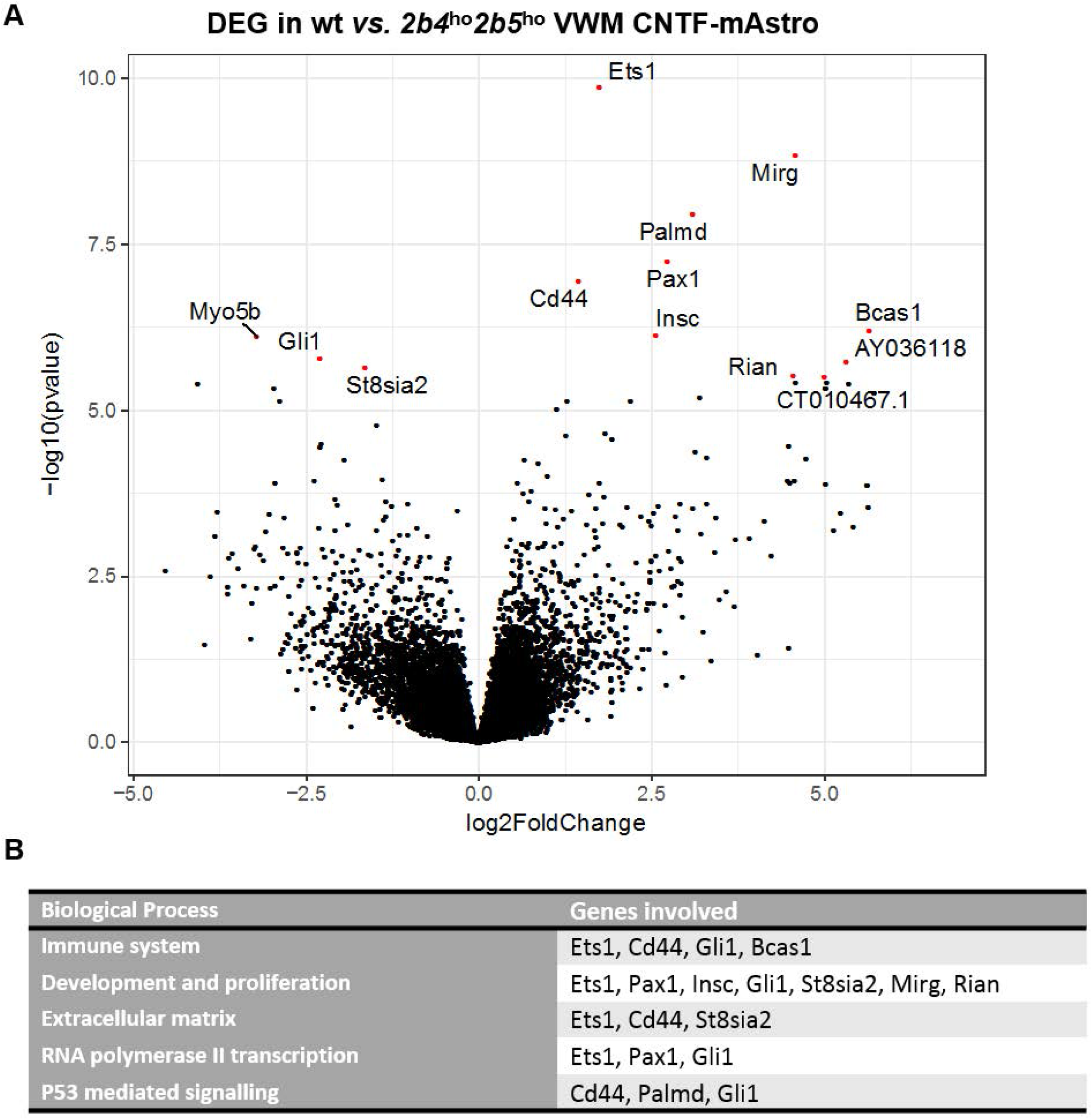
Differentially expressed genes between wt and *2b5*^ho^ CNTF-mAstro. Volcano plot of differential expression analysis between *wt* (n = 4) and *2b4*^ho^*2b5*^ho^ (*n* = 4) CNTF-mAstro with significant DEGs colored in red and labeled (A). DEGs shown per significantly enriched GO term (B).

### HiPSCs differentiate into distinctive astrocytic subpopulations

In order to show subtype specific abnormalities in human VWM cells, we generated CNTF-induced human astrocytes (CNTF-hAstro) and FBS-induced human astrocytes (FBS-hAstro) from control iPSC lines. Both subtypes showed immunostaining for astrocyte-associated markers GFAP, Nestin, and S100β (**Fig. 8A-F & Supplementary Fig. 1C-E**). Similar to mAstro cultures, CNTF-hAstro cultures generated cells with many and thin protrusions (**Fig. 8A-C**), while FBS-hAstro cultures showed a more homogeneous population of flat and round astrocytes (**Fig. 8D-H**). Morphological analysis confirmed differences, although they did not reach statistical significance. FBS-hAstro were rounder (**Fig. 8G**; two sample *t*(4) = −1.385, *p* = .238), the surface of FBS-hAstro was larger (**Fig. 8H**; two samples *t*(4)= −1.813, *p* = .144), and the perimeter corrected for total surface area of the cell was higher in CNTF-hAstro compared to FBS-hAstro (**Fig. 8I**; two sample *t*(4) = 0.95, *p* = .*40*). Expression of astrocyte-associated genes was studied by qPCR analysis and showed that all markers were higher expressed in CNTF-hAstro, reaching statistical significance for *SOX9* (2.54 fold change, paired *t*(3) = −2.84, *p* = .026), *Nestin* (2.21 fold change, paired *t*(3) = −3.70, *p* = .034) and *BLBP* (14.07 fold change, paired *t*(3) = −4.1, *p* = .026) as compared to FBS-hAstro (**Fig. 8J**). To investigate whether the two hAstro subtypes could show a reactivity response, cultures were treated with polyinosinic:polycytidylic acid (Poly(I:C)), a synthetic dsRNA and ligand to TLR3-receptors (Marshall-Clarke *et al*., 2007). mRNA expression levels of a number of genes involved in the astrocytic stress response (Clarke *et al*., 2018) were evaluated by qPCR. A comparison of the expression of these genes in the vehicle condition revealed no differences between subtypes and/or genotypes, except for *FBLN5* (**Supplementary Fig. 1F**, one-way ANOVA *F*(3,10) = 5.38, *p* = .02, post-hoc Dunnett Control CNTF-hAstro *vs*. Control FBS-hAstro *p* = .023, VWM CNTF-hAstro *vs*. Control FBS-hAstro *p* = .015). Both CNTF-hAstros and FBS-hAstros showed significant upregulation of *TLR3* after Poly(I:C)-treatment as compared to cells in the H_2_O-treated control condition (**Fig. 8K**; CNTF-hAstros: paired *t*(3) = 3.62, *p* = .036; **Fig. 8L**, FBS-hAstros: paired *t*(3) = 5.01, *p* = .015). All reactivity-related genes that were tested to assess downstream stress-related effects of TLR3 activation were expressed higher by Poly(I:C)-treated hAstro. After Poly(I:C) treatment, CNTF-hAstro showed a significant upregulation in *FKBP5* (paired *t*(3) = 3.38, *p* = .043) and *PTX3* (**Fig. 8K**; paired *t*(3) = 4.44, *p* = .021). FBS-hAstro showed a significant increase in *FKBP5* (**Fig. 8L**; paired *t*(3) = 3.45, *p* = .041) and a trend in *PTX3* (paired *t*(3) = 3.11, *p* = .053). Both CNTF-hAstro and FBS-hAstro showed a trend for upregulation of *SERPING1* (paired *t*(3) = 2.96, *p* = .060 and paired *t*(3) = 2.92, *p* = .061, respectively). In short, both CNTF- and FBS-hAstro showed appropriate upregulation of reactivity-related genes in response to a cellular stressor, but no differences in reactive response between control CNTF-hAstro and FBS-hAstro were observed.

**Fig. 8.**
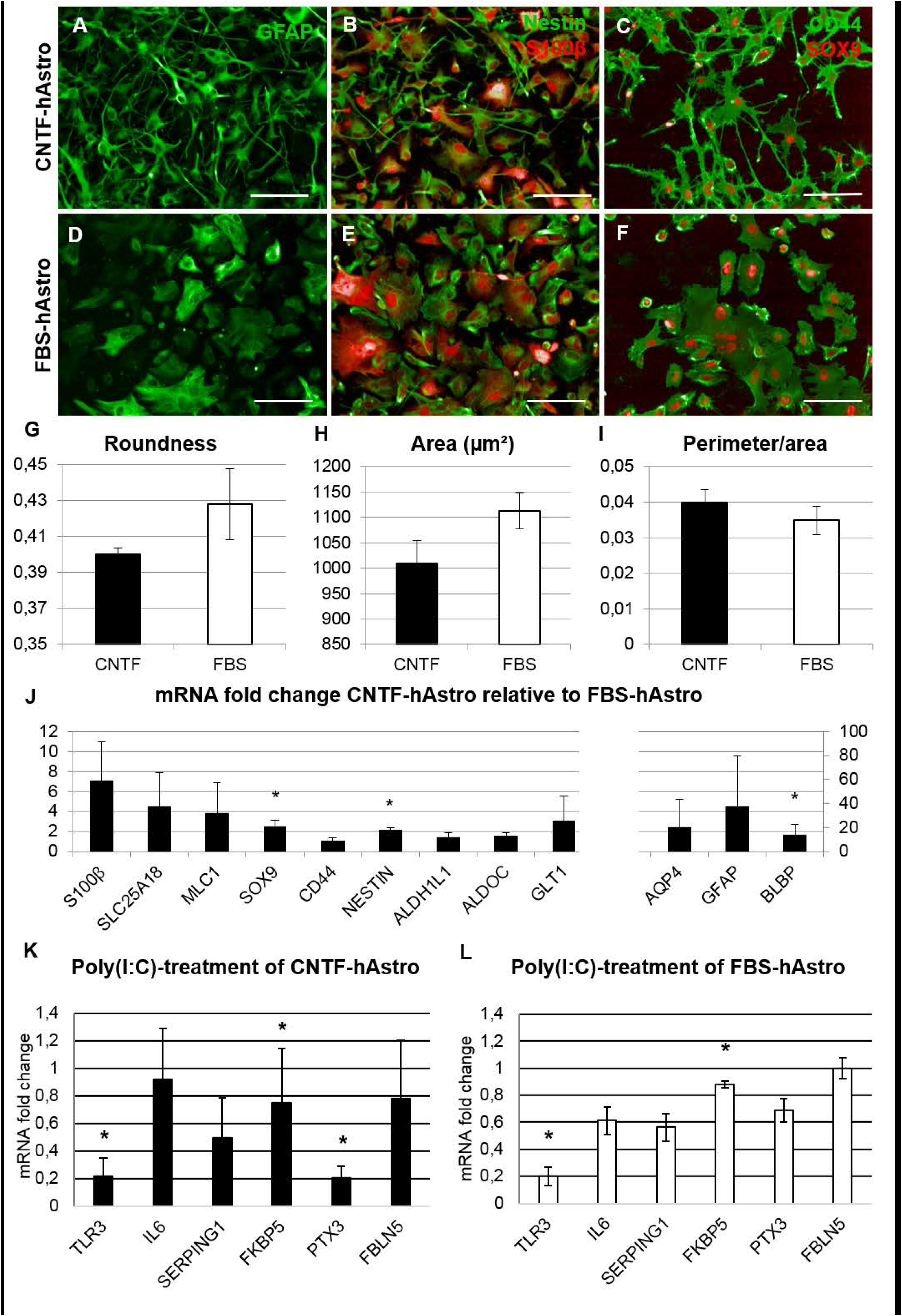
Control CNTF- and FBS-hAstro subtypes differ in morphology and mRNA expression level. Immunostaining showed expression of astrocyte markers GFAP (A&D), Nestin and S100B (B&E), and CD44 and SOX9 (C&F) in both control CNTF-hAstro (B–D) and FBS-hAstro (E–G). Scalebar = 100 μm. Morphological analysis showed differences in roundness (G), surface area (H), and perimeter corrected for area (I) between subpopulations of CNTF-hAstro (*n* = 7) and FBS-hAstro (*n* = 6) based on CD44 immunostaining. A qPCR on control CNTF-hAstro (*n* = 4) showed differential expression of *SOX9*, and *BLBP* relative to FBS-hAstro (*n* = 4). * = *p* < .05 (J). Treatment with Poly(I:C) elicited an upregulation of mRNA expression levels of all tested reactivity-associated genes as well as *TLR3* in both CNTF-hAstro (*n* = 8; K) and FBS-hAstro (*n* = 8; L) as compared to the vehicle (dH2O)condition. CNTF-hAstro showed a significant fold change increase of *TLR3, FKBP5* and *PTX3* mRNA. FBS-hAstro showed a significant increase in *TLR3* and *FKBP5* mRNA. * = *p* <.05. Bars represent mean ± SEM.

### CNTF-hAstro and FBS-hAstro show distinctive gene expression profiles

As, to our knowledge, there are no transcriptome profiles of cultured, or purified, human grey and white matter astrocytes, we performed whole-genome transcriptome analysis on CNTF-hAstro and FBS-hAstro. **Fig. 9A** shows a volcano plot of DEGs in CNTF-hAstro compared to FBS-hAstro. In total, 346 genes were significantly differentially expressed between the subtypes (**Supplementary Table 2**). Enrichment analysis with GO terms (Molecular functions) on the DEGs between CNTF-hAstro and FBS-hAstro could be classified in the categories ‘Receptor/ligand activity’, ‘Extracellular space’, ‘Enzymatic activity’, ‘Immune system’ and ‘Cytoskeleton’ (**Fig. 9B**). These findings show that we generated two human astrocyte subpopulations *in vitro*: white matter-associated CNTF-hAstro and grey matter-associated FBS-hAstro, with specific morphological characteristics and differential expression profiles.

**Fig. 9.**
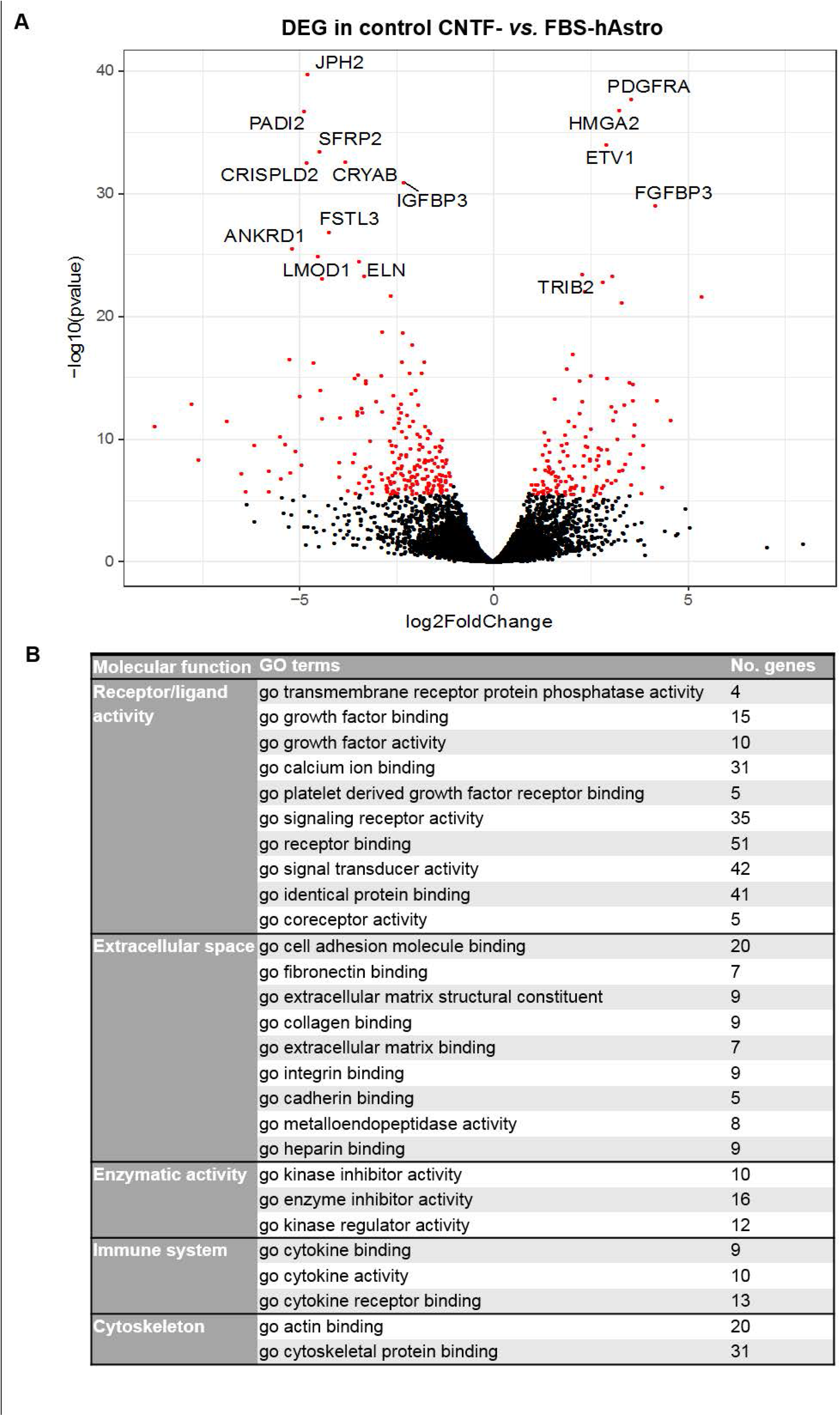
Differentially expressed genes between control CNTF-hAstro and FBS-hAstro. Volcano plot of differential expression analysis between CNTF-hAstro and FBS-hAstro shows significant DEGs in red (A). The 15 most significantly up- or down-regulated DEGs are labeled. DEGs per significantly enriched GO terms (molecular function) were classified in different categories (B).

### VWM mutations predominantly affect CNTF-hAstro over FBS-hAstro

To study astrocyte subtype abnormalities in VWM, we generated CNTF-hAstro and FBS-hAstro populations from control and VWM hiPSCs. To perform unbiased differential expression analysis, we performed transcriptome analysis of VWM and control CNTF-hAstro. In total, 63 genes were significantly differentially expressed between the VWM and control CNTF-hAstro (**Supplementary Table 3**). A volcano plot of covariate-corrected DEGs in VWM compared control CNTF-is shown in **Fig. 10A**. When comparing control and VWM FBS-hAstro, 37 DEGs were detected (**Supplementary Table 3**), which is almost half the number of DEGs found in the CNTF-hAstro subtype, suggesting the FBS-hAstro were less affected by VWM mutations. A volcano plot is presenting the covariate-corrected DEGs in VWM compared to control FBS-hAstro (**Fig. 10B**). Enrichment analysis of GO terms (Biological processes) on the DEGs between control and VWM CNTF-hAstro could be categorized as ‘Immune system’, ‘Extracellular space’, ‘Cell development’, ‘Neuronal functioning’ and ‘Vasculature-related’, whereas the DEGs between control and VWM FBS-hAstro did not show significant enrichment in the latter two categories (**Fig. 10C**). Full lists of all significant GO terms are available upon request. Since these results demonstrate that CNTF-hAstro presented a more broadly modulated transcript profile by a VWM genotype than FBS-hAstro, the differences between VWM and control CNTF-hAstro were further investigated. VWM compared to control CNTF-hAstro showed different morphologies (**Fig. 10D, E**), although these did not reach statistical significance when quantified using automated Columbus^®^ software. The VWM CNTF-hAstro were rounder (**Fig. 10F**; two sample *t*(4) = 0.62, *p* = .58), showed a larger surface area (**Fig. 10G**; two sample *t*(4) = 1.78, *p* = .15), and a reduced perimeter corrected for the surface area (**Fig. 10H**; two sample *t*(4) = −1.00, *p* = .37) compared to control CNTF-hAstro. Furthermore, expression analysis for astrocyte-associated markers by qPCR analysis (Fig. 10I) showed that VWM CNTF-hAstro had a significantly higher expression of *AQP4* (two sample *t*(5) = 2.94, *p* = .032) and *NESTIN* (two sample *t*(5) = 3.61, *p* = .015). These findings demonstrate that CNTF-hAstro were more profoundly affected by the VWM genotype than were FBS-hAstro.

**Fig. 10.**
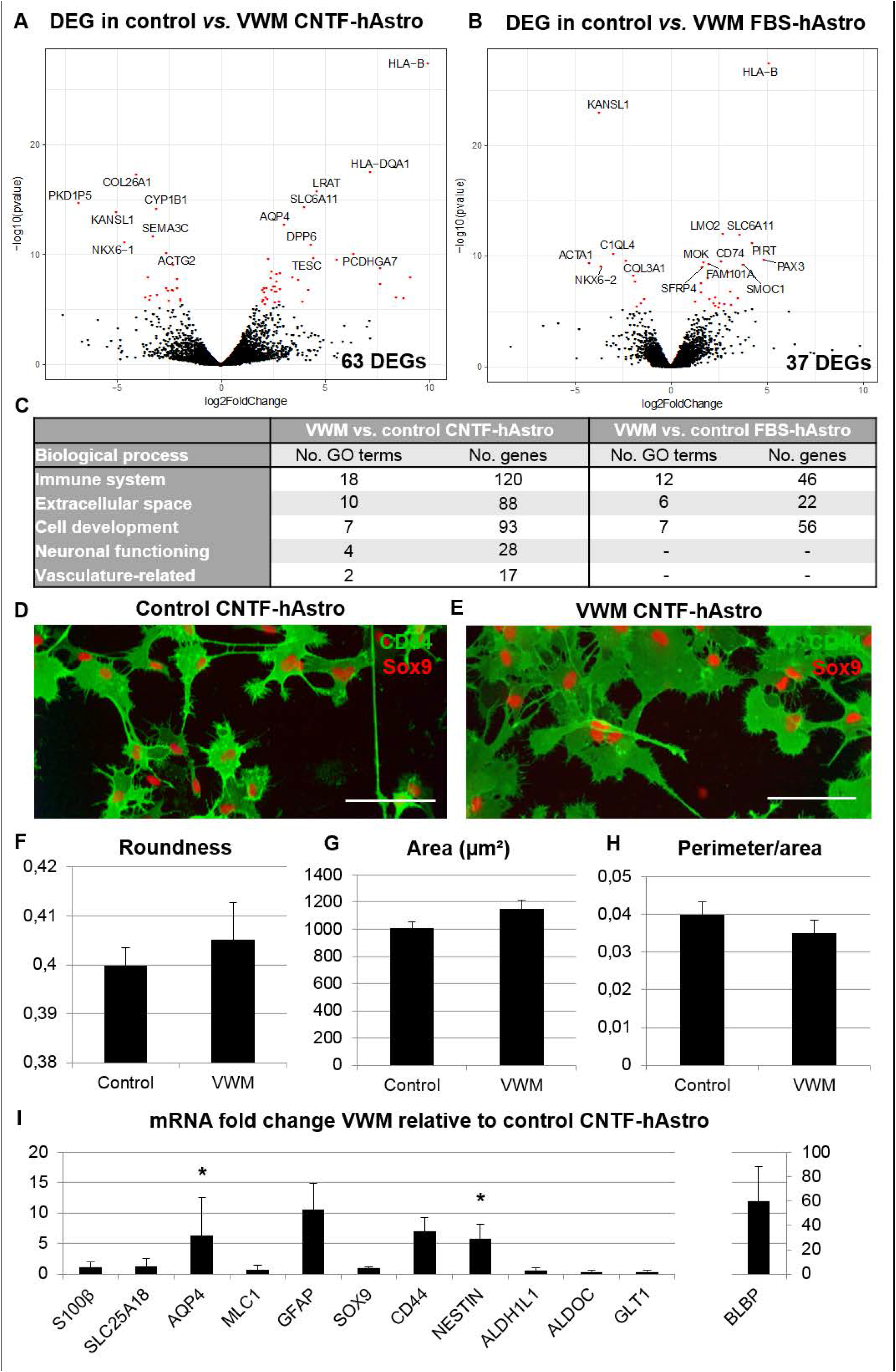
VWM hAstro differ in morphology and mRNA expression level from control hAstro. Differential expression analysis revealed significant DEGs between control and VWM in CNTF-hAstro (control *n* = 3, VWM *n* = 4; A) and FBS-hAstro (control *n* = 4, VWM *n* = 4; B). The volcano plot indicates significant DEGs in red with the top 15 most significantly up- or down-regulated DEGs labeled. DEGs per significant enriched GO terms (biological process) were classified in different categories (C). Morphology of control (D) and VWM (E) CNTF-hAstro was visualised using CD44 and SOX9 immunostaining. Morphological analysis showed differences in roundness (F), area (G), and perimeter corrected for area (H) between VWM and control CNTF-hAstro based on the CD44 immunostaining. A qPCR on VWM CNTF-hAstro (*n* = 4) showed differential expression of *S100B, SLC25A18, MLC1, SOX9, CD44, Nestin, ALDH1L1, AldoC, SLC1A2, AQP4, GFAP* and *BLBP* relative to control CNTF-hAstro (*n* = 4; I). * = *p* < .05. Bars represent mean ± SEM.

## Discussion

In this study we aimed to generate functional astrocytes and specify differentiation protocols for white and grey matter-like astrocyte subtypes from hiPSCs and miPSCs, using respectively CNTF or FBS supplementation. The astrocyte subtypes were characterized for astrocyte-associated markers, typical astrocyte-like morphologies, and reactivity in response to stress. VWM astrocytes showed intrinsic defects. In particular, the white matter-like astrocytes confirmed selective vulnerability for the VWM genotype. Most findings were validated across species, but certain pathways were specifically affected in VWM patient white matter-like astrocytes.

*In vivo*, astrocytes form a heterogeneous group of cells, with many subpopulations that vary in morphology, location and (Bayraktar *et al*., 2014; Molofsky and Deneen, 2015; Tabata, 2015). Functionally grey and white matter astrocytes differ greatly, for example in glutamate uptake (Hassel *et al*., 2003; Goursaud *et al*., 2009) and gap junction coupling (Israel *et al*., 2003; Haas *et al*., 2006; Houades *et al*., 2006). Therefore, we are in need of robust and efficient protocols that generate pure populations of human functional astrocyte subtypes. Even though hiPSC differentiation protocols have been developed to generate astrocytes with varying regional identities (Krencik and Zhang, 2011; Nadadhur *et al*., 2018), so far no protocols to produce white and grey matter astrocytes are in place, which is complicated by the lack of grey and white matter-specific markers. However, increased expression of EGF, FGFR3, GFAP, Nestin, S100B, GLT1, GLAST and CNTF has previously been associated with white matter astrocytes (Pringle *et al*., 2003; Vinukonda *et al*., 2016) (Dallner *et al*., 2002; Hassel *et al*., 2003; Steiner *et al*., 2007; Goursaud *et al*., 2009), as well as a smaller cell size and more complex processes in culture (Raff *et al*., 1983; Hughes *et al*., 1988; Landis *et al*., 1990; Bambrick *et al*., 1996; Haas *et al*., 2012). Here we presented protocols for human and mouse grey and white matter-like astrocytes. Our CNTF-induced astrocytes were smaller, had more complex processes, and showed increased expression of astrocyte markers compared to FBS-induced astrocytes. This shows that these cells are consistent with white and grey matter astrocytes respectively and can be used for *in vitro* disease modeling (Li *et al*., 2018) and for therapy development (Zhang *et al*., 2016a).

The studied markers are not specific for astrocyte subtypes. Increased expression of GFAP, BLBP, Nestin, SOX9 and S100β in astrocytes has been associated with a reactive or immature state (Clarke *et al*., 1994; Van Eldik and Wainwright, 2003; White *et al*., 2010; Sun *et al*., 2017). Several studies suggest that CNTF induces activation of astrocytes (Clatterbuck *et al*., 1996; Wu *et al*., 2006; Seidel *et al*., 2015), for example via the JAK/STAT pathway (Kahn *et al*., 1997). However, under baseline conditions or after treatment with the cellular stressor Poly(I:C) the astrocyte subtypes did not show differences in expression levels of the reactivity-related genes, except for FBLN5. Upregulation of BLBP and Nestin in CNTF-induced astrocytes could indicate an immature phenotype (Zerlin *et al*., 1995; Anthony *et al*., 2004). On the other hand S100β expression was also upregulated in CNTF-induced astrocytes and is thought to label a more mature population (Raponi *et al*., 2007). Analysis of the RNAseq data showed increased LIFR (LIF receptor) transcript levels in our white matter-like astrocytes, which is associated with neural stem cell differentiation into GFAP-positive astrocytes (Koblar *et al*., 1998), and a mature astrocyte state (Bennett *et al*., 2016; Zhang *et al*., 2016b; Clarke *et al*., 2018). In accordance with this, CNTF-induced astrocytes showed upregulation of glutamate signalling-related DEGs GRIK3 and GRIA1, which indicates mature astrocyte functionality. These findings suggest that our CNTF-induced astrocytes do not represent an immature or a chronically reactive state.

Astrocytes play a central role in pathology in VWM (Bugiani *et al*., 2011; Leferink *et al*., 2017). The presented astrocytes derived from miPSCs and hiPSCs recapitulate earlier findings in mouse and human, such as increased proliferation, morphological abnormalities and induction of decreased OPC maturation (Bugiani *et al*., 2011; Dooves *et al*., 2016). Interestingly, white matter-like astrocytes are more vulnerable to VWM mutations than grey matter-like astrocytes, as indicated before in patient tissue (Bugiani *et al*., 2011; Leferink *et al*., 2017). Using mouse cultures we demonstrated that white matter-like astrocytes, but not grey matter-like astrocytes, inhibited OPC maturation. MiPSC- and hiPSC-based models represented species-specific features. Earlier studies indicated that HA is increased in brains of VWM patients (Bugiani *et al*., 2013), but is not a clear determinant in disease phenotypes in primary cultures of VWM mouse cells (Dooves *et al*., 2016). In contrast, but in line with findings in humans, impaired OPC maturation induced by human VWM astrocytes could be rescued by HYAL treatment. This suggests that iPSCs can be used to study intrinsic differences between astrocyte subtypes and to identify shared and unique disease mechanisms between species.

Transcriptome analyses on hiPSC and miPSC cultures both suggest involvement of immune system and extracellular matrix in VWM pathology. In mouse cultures only 13 DEGs between VWM and control were found, of which some genes have been related to VWM disease mechanisms before. *Cd44* encodes a transmembrane receptor that regulates cellular responses to HA, was significantly upregulated in VWM compared to control mouse white matter-like astrocytes. Previous studies showed an increase in CD44 expression in VWM patient white matter astrocytes in post-mortem tissue (Bugiani *et al*., 2013). Further, VWM mouse cultures showed increased *Gli1* expression, which codes for a transcription factor that is a direct target of Sonic Hedgehog (SHH) signaling. SHH regulates proliferation of neural progenitor cells together with intracellular receptor Smoothened (SMO) and has neuroprotective effects (Patel *et al*., 2017). While not significant, *SHH* and *SMO* levels were also decreased (2-2.5 fold) in VWM white matter-like astrocytes. A recent study showed an impaired SHH pathway in primary astrocytes from the *Eif2b5^R132H/R132H^* mouse, confirming the involvement of the SHH pathway in VWM (Atzmon *et al*., 2018). Interestingly, VWM mouse cultures also showed increased *Ets1* expression. *Ets1* is a transcription factor that is involved in the T-cell immune response and was shown to be upregulated in astrocytes surrounding white matter lesions in a mouse model for multiple sclerosis (Gerhauser *et al*., 2007). These findings suggest that the SHH pathway and immune response are interesting targets for VWM.

Exclusively in the human white matter-like astrocytes, differences between VWM and control were associated with the GO terms ‘Neuronal functioning’ and ‘Vasculature-related’. The ‘Neuronal functioning’-related DEGs include upregulation of *GRID1, GRIA2*, and *GRIN2A* in VWM which encode subunits of membrane ionotropic NMDA and AMPA receptors. A downstream effect of activation of these receptors is stimulation of the ATP-driven Na^+^ pump. This induces glycolytic upregulation of ATP and lactate in astrocytes, as well as regulation of the extracellular potassium concentration because the ATP-driven Na+ pump actively pumps potassium into the cells (Lanciotti *et al*., 2013). The channel protein AQP4 is regulated by extracellular potassium levels and involved in water permeability of astrocytes (Nagelhus and Ottersen, 2013), which can lead to neurotoxicity and myelin defects when dysregulated (Plog and Nedergaard, 2018). Interestingly, VWM white matter-like astrocytes showed increased AQP4 levels. DEG analysis includes increased expression of *SLC6A1, SLC6A11*, and *GABBR2* in VWM, which encode major transmembrane GABA transporter GAT-1, GAT-3, and GABAb receptor subunit 2, respectively. Increased extracellular levels of GABA, released by interneurons for example, lead to astrocytic Ca2^+^ elevations mediated by GABAb receptor activation (Guerra-Gomes *et al*., 2017). This may further lead to astrocytic glutamate release, thereby potentiating inhibitory synaptic transmission (Kang *et al*., 1998), and to astrocytic ATP/adenosine-mediated heterosynaptic suppression affecting excitatory transmission (Boddum *et al*., 2016). So, the transcript analysis provided new insights into human specific signalling pathways possibly involved in VWM pathophysiology.

In conclusion, we have robustly created and characterized white and grey matter-like astrocyte subtypes using both mouse and human iPSCs. These specific subtypes are highly important to study diseases where specific astrocytes are affected, and for cellular replacement of those cells. By combining human and mouse models, we confirmed the selective and intrinsic vulnerability of white matter-like astrocytes for VWM mutations, and suggest involvement of the extracellular matrix and immune system in VWM disease mechanisms. The RNAseq profiles can serve as reference databases for validation and characterization of astrocyte subtype in both human and mouse. Future studies can benefit from new insights in VWM, and can apply developed protocols for the generation of subtype specific astrocytes in disease modeling and drug testing studies.

## Acknowledgements

We would like to thank Jurjen Broeke for his help with the calcium imaging of the astrocytes. We would like to thank Aina Badia for her help with the morphological analysis using Columbus^®^ software.

This research is funded by a ZonMw VIDI research grant (91712343; Vivi M. Heine), an European Leukodystrophy Association (ELA) Research Grant (2014-012L1; Vivi M. Heine), an E-Rare Joint Call project (9003037601; Vivi M. Heine), and an NWO Spinoza grant (Marjo S. van der Knaap).

The authors declare no conflict of interest.

**Supplementary material**

## Abbreviations

ACM: astrocyte-conditioned medium
CNS: central nervous system
CNTF: ciliary neurotrophic factor
DEG: Differentially expressed gene
EB: embryoid body
EGF: epidermal growth factor
FBS: Fetal bovine serum
FGF: fibroblast growth factor
GO: Gene ontology
GSA: Global Screening Array
HA: hyaluronic acid/ hyaluronan
HYAL: Hyaluronidase
hiPSCs: human induced pluripotent stem cells
hAstro: human iPSC-derived astrocytes
iPSC: induced pluripotent stem cells
mAstro: mouse iPSC-derived astrocytes
mESC: mouse embryonic stem cells
miPSC: mouse induced pluripotent stem cells
OPC: oligodendrocyte precursor cell
Poly(I:C): polyinosinic:polycytidylic acid
RA: Retinoic acid
RI: Rock inhibitor
RNAseq: RNA sequencing
VWM: Vanishing White Matter
Wt: wild type

## References

Anthony TE, Klein C, Fishell G, Heintz N. Radial glia serve as neuronal progenitors in all regions of the central nervous system. Neuron 2004; 41(6): 881–90.

Atzmon A, Herrero M, Sharet-Eshed R, Gilad Y, Senderowitz H, Elroy-Stein O. Drug Screening Identifies Sigma-1-Receptor as a Target for the Therapy of VWM Leukodystrophy. Front Mol Neurosci 2018; 11: 336.

Back SA, Tuohy TM, Chen H, Wallingford N, Craig A, Struve J, et al. Hyaluronan accumulates in demyelinated lesions and inhibits oligodendrocyte progenitor maturation. Nat Med 2005; 11(9): 966–72.

Bambrick LL, de Grip A, Seenivasan V, Krueger BK, Yarowsky PJ. Expression of glial antigens in mouse astrocytes: species differences and regulation in vitro. J Neurosci Res 1996; 46(3): 305–15.

Bayraktar OA, Fuentealba LC, Alvarez-Buylla A, Rowitch DH. Astrocyte development and heterogeneity. Cold Spring Harb Perspect Biol 2014; 7(1): a020362.

Bennett ML, Bennett FC, Liddelow SA, Ajami B, Zamanian JL, Fernhoff NB, et al. New tools for studying microglia in the mouse and human CNS. Proc Natl Acad Sci U S A 2016; 113(12): E1738–46.

Boddum K, Jensen TP, Magloire V, Kristiansen U, Rusakov DA, Pavlov I, et al. Astrocytic GABA transporter activity modulates excitatory neurotransmission. Nat Commun 2016; 7: 13572.

Bugiani M, Boor I, van Kollenburg B, Postma N, Polder E, van Berkel C, et al. Defective glial maturation in vanishing white matter disease. J Neuropathol Exp Neurol 2011; 70(1): 69–82.

Bugiani M, Postma N, Polder E, Dieleman N, Scheffer PG, Sim FJ, et al. Hyaluronan accumulation and arrested oligodendrocyte progenitor maturation in vanishing white matter disease. Brain 2013; 136(Pt 1): 209–22.

Chandrasekaran A, Avci HX, Leist M, Kobolak J, Dinnyes A. Astrocyte Differentiation of Human Pluripotent Stem Cells: New Tools for Neurological Disorder Research. Front Cell Neurosci 2016; 10: 215.

Clarke LE, Liddelow SA, Chakraborty C, Munch AE, Heiman M, Barres BA. Normal aging induces A1-like astrocyte reactivity. Proc Natl Acad Sci U S A 2018; 115(8): E1896–E905.

Clarke SR, Shetty AK, Bradley JL, Turner DA. Reactive astrocytes express the embryonic intermediate neurofilament nestin. Neuroreport 1994; 5(15): 1885–8.

Clatterbuck RE, Price DL, Koliatsos VE. Ciliary neurotrophic factor stimulates the expression of glial fibrillary acidic protein by brain astrocytes in vivo. J Comp Neurol 1996; 369(4): 543–51.

Dallner C, Woods AG, Deller T, Kirsch M, Hofmann HD. CNTF and CNTF receptor alpha are constitutively expressed by astrocytes in the mouse brain. Glia 2002; 37(4): 374–8.

Das PP, Hendrix DA, Apostolou E, Buchner AH, Canver MC, Beyaz S, et al. PRC2 Is Required to Maintain Expression of the Maternal Gtl2-Rian-Mirg Locus by Preventing De Novo DNA Methylation in Mouse Embryonic Stem Cells. Cell Rep 2015; 12(9): 1456–70.

Dashzeveg N, Taira N, Lu ZG, Kimura J, Yoshida K. Palmdelphin, a novel target of p53 with Ser46 phosphorylation, controls cell death in response to DNA damage. Cell Death Dis 2014; 5: e1221.

Diaz-Amarilla P, Olivera-Bravo S, Trias E, Cragnolini A, Martinez-Palma L, Cassina P, et al. Phenotypically aberrant astrocytes that promote motoneuron damage in a model of inherited amyotrophic lateral sclerosis. Proc Natl Acad Sci U S A 2011; 108(44): 18126–31.

Dooves S, Bugiani M, Postma NL, Polder E, Land N, Horan ST, et al. Astrocytes are central in the pathomechanisms of vanishing white matter. J Clin Invest 2016; 126(4): 1512–24.

Emdad L, D’Souza SL, Kothari HP, Qadeer ZA, Germano IM. Efficient differentiation of human embryonic and induced pluripotent stem cells into functional astrocytes. Stem Cells Dev 2012; 21(3): 404–10.

Falk A, Heine VM, Harwood AJ, Sullivan PF, Peitz M, Brustle O, et al. Modeling psychiatric disorders: from genomic findings to cellular phenotypes. Mol Psychiatry 2016; 21(9): 1321.

Gerhauser I, Alldinger S, Baumgartner W. Ets-1 represents a pivotal transcription factor for viral clearance, inflammation, and demyelination in a mouse model of multiple sclerosis. J Neuroimmunol 2007; 188(1-2): 86–94.

Goursaud S, Kozlova EN, Maloteaux JM, Hermans E. Cultured astrocytes derived from corpus callosum or cortical grey matter show distinct glutamate handling properties. J Neurochem 2009; 108(6): 1442–52.

Guerra-Gomes S, Sousa N, Pinto L, Oliveira JF. Functional Roles of Astrocyte Calcium Elevations: From Synapses to Behavior. Front Cell Neurosci 2017; 11: 427.

Gupta K, Chandran S, Hardingham GE. Human stem cell-derived astrocytes and their application to studying Nrf2-mediated neuroprotective pathways and therapeutics in neurodegeneration. Br J Clin Pharmacol 2013; 75(4): 907–18.

Haas B, Schipke CG, Peters O, Sohl G, Willecke K, Kettenmann H. Activity-dependent ATP-waves in the mouse neocortex are independent from astrocytic calcium waves. Cereb Cortex 2006; 16(2): 237–46.

Haas C, Neuhuber B, Yamagami T, Rao M, Fischer I. Phenotypic analysis of astrocytes derived from glial restricted precursors and their impact on axon regeneration. Exp Neurol 2012; 233(2): 717–32.

Han Z, He H, Zhang F, Huang Z, Liu Z, Jiang H, et al. Spatiotemporal expression pattern of Mirg, an imprinted non-coding gene, during mouse embryogenesis. J Mol Histol 2012; 43(1): 1–8.

Hassel B, Boldingh KA, Narvesen C, Iversen EG, Skrede KK. Glutamate transport, glutamine synthetase and phosphate-activated glutaminase in rat CNS white matter. A quantitative study. J Neurochem 2003; 87(1): 230–7.

Houades V, Rouach N, Ezan P, Kirchhoff F, Koulakoff A, Giaume C. Shapes of astrocyte networks in the juvenile brain. Neuron Glia Biol 2006; 2(1): 3–14.

Hughes SM, Lillien LE, Raff MC, Rohrer H, Sendtner M. Ciliary neurotrophic factor induces type-2 astrocyte differentiation in culture. Nature 1988; 335(6185): 70–3.

Ishimoto T, Ninomiya K, Inoue R, Koike M, Uchiyama Y, Mori H. Mice lacking BCAS1, a novel myelin-associated protein, display hypomyelination, schizophrenia-like abnormal behaviors, and upregulation of inflammatory genes in the brain. Glia 2017; 65(5): 727–39.

Israel JM, Schipke CG, Ohlemeyer C, Theodosis DT, Kettenmann H. GABAA receptor-expressing astrocytes in the supraoptic nucleus lack glutamate uptake and receptor currents. Glia 2003; 44(2): 102–10.

Kahn MA, Huang CJ, Caruso A, Barresi V, Nazarian R, Condorelli DF, et al. Ciliary neurotrophic factor activates JAK/Stat signal transduction cascade and induces transcriptional expression of glial fibrillary acidic protein in glial cells. J Neurochem 1997; 68(4): 1413–23.

Kang J, Jiang L, Goldman SA, Nedergaard M. Astrocyte-mediated potentiation of inhibitory synaptic transmission. Nat Neurosci 1998; 1(8): 683–92.

Kleiderman S, Sa JV, Teixeira AP, Brito C, Gutbier S, Evje LG, et al. Functional and phenotypic differences of pure populations of stem cell-derived astrocytes and neuronal precursor cells. Glia 2016; 64(5): 695–715.

Koblar SA, Turnley AM, Classon BJ, Reid KL, Ware CB, Cheema SS, et al. Neural precursor differentiation into astrocytes requires signaling through the leukemia inhibitory factor receptor. Proc Natl Acad Sci U S A 1998; 95(6): 3178–81.

Krencik R, Zhang SC. Directed differentiation of functional astroglial subtypes from human pluripotent stem cells. Nat Protoc 2011; 6(11): 1710–7.

Kuegler PB, Baumann BA, Zimmer B, Keller S, Marx A, Kadereit S, et al. GFAP-independent inflammatory competence and trophic functions of astrocytes generated from murine embryonic stem cells. Glia 2012; 60(2): 218–28.

Lanciotti A, Brignone MS, Bertini E, Petrucci TC, Aloisi F, Ambrosini E. Astrocytes: Emerging Stars in Leukodystrophy Pathogenesis. Transl Neurosci 2013; 4(2).

Landis DM, Weinstein LA, Skordeles CJ. Serum influences the differentiation of membrane structure in cultured astrocytes. Glia 1990; 3(3): 212–21.

Leferink PS, Breeuwsma N, Bugiani M, van der Knaap MS, Heine VM. Affected astrocytes in the spinal cord of the leukodystrophy vanishing white matter. Glia 2017.

Leung C, Jia Z. Mouse Genetic Models of Human Brain Disorders. Front Genet 2016; 7: 40.

Li L, Tian E, Chen X, Chao J, Klein J, Qu Q, et al. GFAP Mutations in Astrocytes Impair Oligodendrocyte Progenitor Proliferation and Myelination in an hiPSC Model of Alexander Disease. Cell Stem Cell 2018; 23(2): 239–51 e6.

Lundgaard I, Osorio MJ, Kress BT, Sanggaard S, Nedergaard M. White matter astrocytes in health and disease. Neuroscience 2014; 276: 161–73.

Marshall-Clarke S, Downes JE, Haga IR, Bowie AG, Borrow P, Pennock JL, et al. Polyinosinic acid is a ligand for toll-like receptor 3. J Biol Chem 2007; 282(34): 24759–66.

Martinian L, Boer K, Middeldorp J, Hol EM, Sisodiya SM, Squier W, et al. Expression patterns of glial fibrillary acidic protein (GFAP)-delta in epilepsy-associated lesional pathologies. Neuropathol Appl Neurobiol 2009; 35(4): 394–405.

Middeldorp J, van den Berge SA, Aronica E, Speijer D, Hol EM. Specific human astrocyte subtype revealed by affinity purified GFAP antibody; unpurified serum cross-reacts with neurofilament-L in Alzheimer. PLoS One 2009; 4(11): e7663.

Molofsky AV, Deneen B. Astrocyte development: A Guide for the Perplexed. Glia 2015; 63(8): 1320–9.

Nadadhur AG, Leferink PS, Holmes D, Hinz L, Cornelissen-Steijger P, Gasparotto L, et al. Patterning factors during neural progenitor induction determine regional identity and differentiation potential in vitro. Stem Cell Res 2018; 32: 25–34.

Nagelhus EA, Ottersen OP. Physiological roles of aquaporin-4 in brain. Physiol Rev 2013; 93(4): 1543–62.

Oksanen M, Petersen AJ, Naumenko N, Puttonen K, Lehtonen S, Gubert Olive M, et al. PSEN1 Mutant iPSC-Derived Model Reveals Severe Astrocyte Pathology in Alzheimer’s Disease. Stem Cell Reports 2017; 9(6): 1885–97.

Patel SS, Tomar S, Sharma D, Mahindroo N, Udayabanu M. Targeting sonic hedgehog signaling in neurological disorders. Neurosci Biobehav Rev 2017; 74(Pt A): 76–97.

Plog BA, Nedergaard M. The Glymphatic System in Central Nervous System Health and Disease: Past, Present, and Future. Annu Rev Pathol 2018; 13: 379–94.

Pringle NP, Yu WP, Howell M, Colvin JS, Ornitz DM, Richardson WD. Fgfr3 expression by astrocytes and their precursors: evidence that astrocytes and oligodendrocytes originate in distinct neuroepithelial domains. Development 2003; 130(1): 93–102.

Raff MC, Abney ER, Cohen J, Lindsay R, Noble M. Two types of astrocytes in cultures of developing rat white matter: differences in morphology, surface gangliosides, and growth characteristics. J Neurosci 1983; 3(6): 1289–300.

Rakela B, Brehm P, Mandel G. Astrocytic modulation of excitatory synaptic signaling in a mouse model of Rett syndrome. Elife 2018; 7.

Raponi E, Agenes F, Delphin C, Assard N, Baudier J, Legraverend C, et al. S100B expression defines a state in which GFAP-expressing cells lose their neural stem cell potential and acquire a more mature developmental stage. Glia 2007; 55(2): 165–77.

Russo FB, Freitas BC, Pignatari GC, Fernandes IR, Sebat J, Muotri AR, et al. Modeling the Interplay Between Neurons and Astrocytes in Autism Using Human Induced Pluripotent Stem Cells. Biol Psychiatry 2018; 83(7): 569–78.

Seidel JL, Faideau M, Aiba I, Pannasch U, Escartin C, Rouach N, et al. Ciliary neurotrophic factor (CNTF) activation of astrocytes decreases spreading depolarization susceptibility and increases potassium clearance. Glia 2015; 63(1): 91–103.

Shaltouki A, Peng J, Liu Q, Rao MS, Zeng X. Efficient generation of astrocytes from human pluripotent stem cells in defined conditions. Stem Cells 2013; 31(5): 941–52.

Sloane JA, Batt C, Ma Y, Harris ZM, Trapp B, Vartanian T. Hyaluronan blocks oligodendrocyte progenitor maturation and remyelination through TLR2. Proc Natl Acad Sci U S A 2010; 107(25): 11555–60.

Steiner J, Bernstein HG, Bielau H, Berndt A, Brisch R, Mawrin C, et al. Evidence for a wide extra-astrocytic distribution of S100B in human brain. BMC Neurosci 2007; 8: 2.

Sun W, Cornwell A, Li J, Peng S, Osorio MJ, Aalling N, et al. SOX9 Is an Astrocyte-Specific Nuclear Marker in the Adult Brain Outside the Neurogenic Regions. J Neurosci 2017; 37(17): 4493–507.

Tabata H. Diverse subtypes of astrocytes and their development during corticogenesis. Front Neurosci 2015; 9: 114.

Takahashi K, Tanabe K, Ohnuki M, Narita M, Ichisaka T, Tomoda K, et al. Induction of pluripotent stem cells from adult human fibroblasts by defined factors. Cell 2007; 131(5): 861–72.

Tcw J, Wang M, Pimenova AA, Bowles KR, Hartley BJ, Lacin E, et al. An Efficient Platform for Astrocyte Differentiation from Human Induced Pluripotent Stem Cells. Stem Cell Reports 2017; 9(2): 600–14.

Van der Knaap MS, Wolf, N.I., Heine, V.M. Leukodystrophies: Five new things. Neurology: Clinical Practice 2016; August 23.

Van Eldik LJ, Wainwright MS. The Janus face of glial-derived S100B: beneficial and detrimental functions in the brain. Restor Neurol Neurosci 2003; 21(3-4): 97–108.

Vinukonda G, Hu F, Mehdizadeh R, Dohare P, Kidwai A, Juneja A, et al. Epidermal growth factor preserves myelin and promotes astrogliosis after intraventricular hemorrhage. Glia 2016; 64(11): 1987–2004.

White RE, McTigue DM, Jakeman LB. Regional heterogeneity in astrocyte responses following contusive spinal cord injury in mice. J Comp Neurol 2010; 518(8): 1370–90.

Williams EC, Zhong X, Mohamed A, Li R, Liu Y, Dong Q, et al. Mutant astrocytes differentiated from Rett syndrome patients-specific iPSCs have adverse effects on wild-type neurons. Hum Mol Genet 2014; 23(11): 2968–80.

Wu Y, Liu RG, Zhou JP. Effect of ciliary neurotrophic factor on activation of astrocytes in vitro. Neurosci Bull 2006; 22(6): 315–22.

Yamanaka K, Komine O. The multi-dimensional roles of astrocytes in ALS. Neurosci Res 2018; 126: 31–8.

Zerlin M, Levison SW, Goldman JE. Early patterns of migration, morphogenesis, and intermediate filament expression of subventricular zone cells in the postnatal rat forebrain. J Neurosci 1995; 15(11): 7238–49.

Zhang K, Chen C, Yang Z, He W, Liao X, Ma Q, et al. Sensory Response of Transplanted Astrocytes in Adult Mammalian Cortex In Vivo. Cereb Cortex 2016a; 26(9): 3690–704.

Zhang Y, Sloan SA, Clarke LE, Caneda C, Plaza CA, Blumenthal PD, et al. Purification and Characterization of Progenitor and Mature Human Astrocytes Reveals Transcriptional and Functional Differences with Mouse. Neuron 2016b; 89(1): 37–53.

