## Supplemental Materials and Methods for "Human and mouse iPSC-derived astrocyte subtypes reveal vulnerability in Vanishing White Matter"

### Supplementary information

#### Figure legends supplementary figures and tables

**Supplementary Figure 1**. VWM and control hAstro express astrocyte markers and are functional. mRNA expression of astrocyte markers was assessed using RT-PCR on both control and VWM hAstro (A). Calcium imaging analysis of control and VWM hAstro showed incidental spontaneous activity, and increased intensity of intracellular calcium staining upon a glutamate puff (arrows), followed by recovery to baseline (B). All graphs and gels show a representative example of one control and one VWM hAstro sample. Morphological analysis of control CNTF- and FBS-hAstro. Morphological analysis showed differences in intensity of immunostaining for Nestin (C), S100β (D), GFAP (E) between control CNTF- and FBS-hAstro. Upon qPCR testing of CNTF- and FBS-hAstro for reactivity-associated genes, it was found that CNTF-hAstro showed a significantly higher expression of *FLBN5* mRNA in vehicle conditions as compared to control FBS-hAstro, but not VWM FBS-hAstro (F).

**Supplemental Table 1**. Differential expressed genes between wt vs. $2b4^{ho}2b5^{ho}$ CNTF-mAstro.

**Supplemental Table 2**. Differential expressed genes between control FBS- vs. CNTF-hAstro.

**Supplemental Table 3**. Differential expressed genes between control vs. VWM CNTF-hAstro and between control vs. VWM FBS-hAstro.

#### Supplementary materials & methods

Mouse iPSC production & maintenance

Mouse iPSCs (miPSC) were produced from fibroblasts. Fibroblasts were isolated from finely cut skin pieces by culturing the skin piece at 37°C with 5% $CO_2$ in Fibroblast medium (DMEM + 10% fetal bovine serum (FBS) + 1% L-glutamine + 2% Penicillin/Streptomycin (Pen/Strep) + 1% non-essential amino acids (NEAA) + 1% Sodium Pyruvate + 0.2 μM β-mercaptoethanol) until cells were migrating out. When fibroblast were confluent they were split 1:3 by incubation in Trypsin/EDTA at 37ºC for 5 min. For iPSC generation fibroblasts were plated 4 x $10^4$ cells/well on a 0.1% gelatin coated 6-wells plate one day prior to infection with lentiviral vectors containing the 'Yamanaka factors' Oct4, Klf4, Sox2 and cMyc (Takahashi and Yamanaka, 2006). Transduction was done in infection medium (DMEM + 10% FBS + 0.2 μM β-mercaptoethanol) supplemented with 10 μg/ml polybrene. The day after transduction the virus was aspirated and cells were cultured in fibroblast medium for 6 days. After that medium was switched to mouse ES medium (DMEM/F12 + 10% knockout serum replacement + 1% L-glutamine + 2% Pen/Strep + 1% NEAA + 0.35 μM β-mercaptoethanol). When colonies started to appear, colonies were picked with a 10 μl pipette and plated on a feeder layer of mouse embryonic fibroblasts (MEF). All miPSC lines used in this study were initially produced on a MEF feeder layer, and later switched to the feeder-free 2i system.

miPSCs were maintained in 2i medium (49% DMEM/F12 + 48% Neurobasal + 11 mM Hepes + 0.5x N2 supplement + 0.5x B27 supplement + 25 μg/ml BSA + 1% glutamax + 1% Pen/Strep + 0.2% β-mercaptoethanol) supplemented with 1 x $10^3$ units LIF (Millipore), 3 μM CHIR99021 (Systems Biosciences) and 1 μM PD0325901 (Selleckchem) on 0.1% gelatin coated plates. Colonies were split every 3-4 days with Accutase (Sigma-Aldrich) incubation and plated on a new gelatin plate.

Primary mouse cell isolation

Primary mouse astrocytes and OPCs were isolated from embryonic day 18 (E18) C57/Bl6J wt mice as previously described (Dooves *et al.*, 2016). All mouse procedures were carried out according to the guidelines of the Animal Approval Committee of the VU University Amsterdam. Primary mouse astrocytes were co-cultured with miPSC mixed glial cells in M41 medium for one week. Primary OPCs were cultured in hAstro or mAstro derived astrocyte conditioned medium (ACM) for one week with daily medium refreshment. After 1 week cells were fixed with 2% PFA for 20 min and analyzed by immunostaining (see below).

Human iPSC production & maintenance

Human iPSCs (hIPSCs) were generated from patient and control fibroblasts by using a lentiviral construct expressing the 'Yamanaka factors' coupled to a tomato red tag (Warlich *et al.*, 2011). In short, the reprogramming lentivirus was produced in HEK293T cells using FuGENE 6 (Promega Corporation) in Opti-MEM medium (Thermo Fisher Scientific) for 48 hours, after which the collected virus was concentrated using ultracentrifugation. Fibroblast were seeded in fibroblast medium (15% Fetal Bovine Serum, 1% Non-Essential Amino Acids, 1% Penicillin/Streptomycin (100 U/ml), 0.1% µl β-mercaptoethanol in DMEM/F12) in a density of 30.000 to 60.000 cells per well of a 6-wells plate, depending on passage number and growth rate of the fibroblasts. After overnight incubation at 37°C with 5% $CO_2$, the medium was switched to infection medium (2% FBS in DMEM/F12), and 1 tube of concentrated virus (~20 µl) was added per well. The next day, cells were washed twice with PBS, and switched back to fibroblast medium. After overnight incubation, the infected cells were passaged with 0.05% Trypsin/EDTA (Thermo Fischer Scientific) to GelTrex (Life Technologies)-coated 10cm dish in fibroblast medium. The next day, medium was switched to E7 medium (Life Technologies). Between 10 – 30 days after infection, reprogrammed cells formed colonies, which were picked when red fluorescent expression was diminished, and seeded on GelTrex coated plates in E8 medium (Life Technologies) supplemented with 10 µM Rock Inhibitor (RI; y-27632, SelleckChem). Each picked colony was cultured as a separate line. HiPSC lines were maintained on GelTrex in E8 medium, and passaged as small fragments using 0.5mM EDTA (Life Technologies) twice a week.

Characterization of iPSCs

All miPSCs and hiPSCs were characterized for pluripotency using alkaline phosphatase staining, immunocytochemistry, and RT-PCR analysis for expression of pluripotent markers.

To test for alkaline phosphatase activity, cells were washed once with room temperature (RT) PBS, and fixated by incubating with 2% PFA (Electron Microscopy Sciences) at RT for 20 minutes. After washing the cells with PBS, the cells were incubated in an alkaline-dye mixture (a combination of mixture 1 containing 21 µl sodium nitrite (0.1 M) mixed with 21 µl FRV-Alkaline solution, and mixture 2 containing 21 µl Naphtol AS-BI Alkaline solution in 937 µl deionized water) for 5 – 15 minutes at RT protected from light. After incubation the alkaline-dye mixture was removed, and cells were rinsed twice with deionized water for 2 minutes.

To test differentiation potential of miPSCs into all three germ layers, cells were detached from the plate by Accutase incubation and cultured on an anti-adherent plate in mouse ES medium. Within 4 days EBs were formed. To confirm correct differentiation patterns, for each line EBs were collected for PCR analysis or plated on a poly-L-ornithin (PLO)-coated plate for overnight adherence in order to be used for immunostaining. Additionally, miPSCs injected into a mouse brain formed mature teratomas with bone tissue, cartilage and hair follicles visible on hematoxylin-eosin staining.

HiPSC lines were checked for pluripotency using a RNA sequencing pluritest, next to an EB differentiation assay. HiPSCs were karyotyped or tested on the Infinium Global Screening Array (GSA) for DNA abnormalities.

Immunocytochemistry

After washing once with RT PBS, cells were fixated by incubating with 2% PFA (Electron Microscopy Sciences) at RT for 20 minutes. After washing 6 times for 5 minutes with PBS, cells were incubated in blocking solution (PBS with 5% normal goat serum (Gibco), 0.1% BSA and 0.3% Triton® X-100 (Sigma) for 1 hour at RT, to block aspecific binding of the antibodies. Subsequently, blocking buffer was replaced with primary antibodies (Supplementary Table 2) in blocking solution. Antibodies were incubated for 1 hour at RT in the dark, and then at 4 ºC overnight. After 6 more washing steps of 5 minutes with RT PBS, secondary antibodies in blocking buffer were added, and incubated at RT for 2 hours in the dark. Used secondary antibodies were goat-anti mouse Alexa Fluor 488 (1:1000, Invitrogen, A11029), goat-anti rat Alexa Fluor 594 (1:1000, Invitrogen, A11007), or Alexa Fluor goat-anti rabbit 594 (1:1000, Invitrogen, A11012). After washing for 6 times during 5 minutes with PBS, cells were stained for 2 minutes at RT in the dark with 1:1000 DAPI in PBS. Cells were washed one more time with PBS, before slides were embedded in 7 µl Fluoromount G (Southern Biotech). Cells on chamberslides and coverslips were subsequently analyzed at the Leica DM6000B microscope (Leica Microsystems BV, Rijswijk, the Netherlands), cells in the black clear bottom 96wells plate (Corning) were analyzed using the Opera LX HCS instrument (PerkinElmer).

Human iPSC differentiation towards generic astrocytes

HiPSCs were differentiated towards human astrocytes (hAstro) as described previously (Nadadhur et al., 2018). In short, hiPSCs were incubated in EDTA twice before being dissociated in PBS and allowing for gravity separation in a tube. They were then transferred to a poly-2-hydroxyethyl methacrylate (Sigma)-coated anti-adhesive plate to form EBs overnight in N2B27 medium (a 1:1 mixture of N-2 and B-27-containing media; N-2 medium consists of DMEM/F-12 GlutaMAX, 1× N-2, 5 µg/ml insulin, 1 mM L-glutamine, 100 µm non-essential amino acids, 100 µM 2-mercaptoethanol, 50 U/ml penicillin and 50 mg/ml streptomycin. B-27 medium consists of Neurobasal, 1× B-27, 200 mM L-glutamine, 50 U/ml penicillin and 50 mg/ml streptomycin) supplemented with FGF2 (4 ng/ml, Peprotech), EGF (20 ng/ml, Peprotech), T3 (40 ng/ml, Sigma) and 10 µM RI. Half of the medium was refreshed every other day. After two days, RI was replaced with Retinoic Acid (RA; 10 µM, Sigma). At day 10 the EBs were plated on GelTrex-coated plates in N2B27 medium supplemented with only EGF (20 ng/ml) and T3 (40 ng/ml). After 4 days, when the cells had formed rosette-like structures, the cells were passaged using Accutase. At day 18, the medium was switched to N2B27 medium without vitamin A (N2B27-vitA) supplemented with T3 (40 ng/ml) and EGF (20 ng/ml). At day 37, the medium was switched to N2B27-vitA supplemented with T3 (40 ng/ml), EGF (5 ng/ml), FGF2 (5 ng/ml), noggin (50 ng/ml, Peprotech), Vitamin C (50 ug/ml, Sigma) and laminin (1 ug/ml, Sigma) for 5 days, after which EGF and FGF2 were omitted from the medium. For mixed astrocyte differentiation, the medium was switched to astrocyte medium (ScienCell, Sanbio), and passaged for 2 – 20 more times over a time period of 15 – 60 days, depending on the required maturation state of the hAstro.

Mouse iPSC differentiation towards glial progenitor cells

To differentiate miPSCs into glial progenitor cells, first EBs were formed by free floating culture in mouse ES medium. After 4 days medium was switched to mouse neural induction medium (DMEM/F12 + 1% NEAA + 1% Sodium Pyruvate + 3 mg/ml D-glucose + 0.5% Pen/Strep + 50 µg/ml Apo-Transferrin + 20 nM Progesterone + 30 nM Sodium Selenite + 60 µM Putrescine + 5 µg/ml Insulin + 2 µg/ml Heparin) supplemented with 0.2 µM retinoic acid for 1 day, and then 3 days in mouse neural induction medium supplemented with retinoic acid and 1 µM purmorphamine. After the neural induction phase cells were plated on PLO/fibronectin coated plates in mouse neural maintenance medium (DMEM/F12 + 1% NEAA + 1% Sodium Pyruvate + 3 mg/ml D-glucose + 0.5% Pen/Strep + 100 µg/ml Apo-Transferrin + 20 nM Progesterone + 30 nM Sodium Selenite + 60 µM Putrescine + 25 µg/ml Insulin + 2 µg/ml Heparin) supplemented with 20 ng/ml FGF2 for 12 days. Half of the medium was changed daily throughout the differentiation protocol. At day 10 cells were passaged 1:4-1:8 depending on cell density to a new PLO/fibronectin-coated plate.

Isolation and culturing of primary mouse astrocytes.

Primary astrocytes were isolated from the forebrain of E18 mice as described previously (Dooves *et al.*, 2016). Forebrain was dissected and washed in HBSS-, following by a 10 minute incubation in activated papaïn solution (Sigma-Aldrich). Papaïn was inactivated by the addition of serum-containing astrocyte medium (DMEM/F12 + 10% FBS + 1% L-glutamine + 1% Pen/Strep). The cells were centrifuged for 5 minutes at 1200 rpm, resuspended in astrocyte medium and plated in a density of 1 brain per T-25 flask. From one litter, both $2b5^{ho}$ and $2b5^{he}$ were derived. $2b5^{he}$ astrocytes are indicated here as 'wt' (wild type) astrocytes since VWM is a recessive disease and $2b5^{he}$ mice do not show any phenotype. The $2b5^{he}$ astrocytes are therefore considered to function as wt astrocytes. When astrocytes became confluent they were split 1:3-1:5 to a new flask by Trypsin-EDTA incubation for 5 min at 37ºC. At the fourth passage, cells were frozen in 1:1 astrocyte medium:astrocyte freezing solution (DMEM/F12 + 20% FBS + 20% DMSO) in a density of 2 million cells per vial. A week before the start of the co-culture astrocytes are plated on PLO/laminin coated 8-well chamber slides at a density of 100.000 cells/well in astrocyte medium. At the start of the co-culture, astrocyte medium was removed and cells were washed once with PBS.

BrdU incorporation human iPSC-derived astrocytes

For quantification of proliferation, 20.000 human iPSC-derived astrocytes of each line were plated on GelTrex-coated 13mm coverslips in a 24-wells plate in astrocyte medium (ScienCell, Sanbio B.V.). After 4 days, the cells were treated with either 1:1000 DMSO (vehicle) or labeled with 10μM BrdU (10mM in DMSO; Sigma) in astrocyte medium (ScienCell, Sanbio B.V.), and incubated for 2 hours at 37°C. The cells were washed twice with PBS, and fixated with 4% PFA for 15 minutes at RT. The cells were again washed twice with PBS, and incubated with permeabilization buffer (0.1% Triton® X-100 in PBS) for 20 minutes at RT. Subsequently, the cells were incubated for 10 minutes with 1N HCl on ice, followed by 10 minutes incubation with 2N HCl at RT, and 10 minutes with phosphate/citric acid buffer (182 mM Na2HPO4 with 9 mM citric acid in $dH_2O$, pH 6.0) at RT. The cells were washed with permeabilization buffer three times, 2 minutes each. Cells were then incubated with blocking solution (5% normal goat serum, 0.1% BSA, 0.3% Triton® X-100 in PBS) for 30 minutes at RT, followed by overnight incubation with rat anti-BrdU primary antibody (Supplementary Table 2) in blocking buffer at RT. The next day, cells were washed with permeabilization buffer (3 times, 2 minutes each), and incubated with goat-anti rat Alexa Fluor 594 (1:1000, Invitrogen, A11007) secondary antibody in blocking buffer for one hour at RT. After washing 6 times for 5 minutes with PBS, cells were stained with 1:1000 DAPI in PBS for 2 minutes at RT in the dark. Cells were washed one more time with PBS before embedding the slides in 7 μl Fluoromount G.

For quantification, a Leica DM6000B microscope (Leica Microsystems BV, Rijswijk, the Netherlands) was used to take 6 pictures at random locations per coverslip at 10.00x magnification. ImageJ software with the ITCN plugin was used for automated cell count of DAPI- and BrdU-positive nuclei.

Calcium imaging human iPSC-derived astrocytes

For calcium imaging of glutamate uptake in hIPSC-derived astrocytes, 15.000 cells were plated on a GelTrex coated- 18 mm coverslip in a 12-wells plate, and cultured for 7 days. The cells were incubated for 5 minutes with 1:2000 calcium dye Fluor5 (2mM in DMSO, Molecular probes) in astrocyte medium (ScienCell, Sanbio B.V.). After a baseline recording in continuous flowing aCSF, a 15 seconds puff of 10 μM L-Glutamic acid (Sigma, in aCSF) was given to the cell, followed by 3 minutes of recording, a second 10 μM L-Glutamic acid puff, and a final 3 minutes of recording. Per coverslip, 2 regions were measured at a speed of 2 frames per second. Camera settings: 1 pixel = 16 μm, 20x magnification, recordings are in AVI format. Data was analyzed using Fiji Is Image J (FIJI) software. In each region, 12 - 15 regions of interest (ROI) were selected in the cytoplasm of individual cells, and flurorescent intensity (F) was measured. As a background ($F_0$), the average fluorescent intensity of 100 frames of inactivity was used ($F_0 = (\sum F_{1-100})/100$). To correct for this background, change in relative fluorescence versus time for each ROI (change in fluorescence = $(F-F_0)/F_0$) was calculated, and represented in a graph over the frames.

RNA isolation

After washing cells twice with PBS, total RNA was isolated using 750 μl TRIzol reagent (Invitrogen, Carlsbad, CA, USA) per cell sample. After addition of 150 μl chloroform (Sigma-Aldrich) per 750 μl TRIzol, the sample was shaken vigorously for 15 seconds, followed by 10 min incubation at room temperature (RT). A 10-minute centrifugation at 12000 rpm at 4 ºC resulted in a liquid phase separation, from which the aqueous upper phase containing the RNA was transferred to a new vial. Subsequently, RNA was precipitated by adding 350 μl isopropanol (Sigma-Aldrich) to the vial, which was inverted to mix the solution and incubated at room temperature for 10 minutes. It was then centrifuged for another 10 minutes at 12000 rpm 4 ºC. The pellet was washed twice with 1 ml 70% ethanol and air-dried, before re-dissolving the RNA in 10 μl RNAse free water. Concentrations and purity of RNA were established using a Nanodrop (Thermo Scientific). From 1 μg RNA in 12.2 μl, cDNA was synthesized. To each sample 4 μl 5x First strand buffer (Invitrogen), 1 μl Random hexamer oligo (Qiagen) (0,6μg/μl), 1 μl Oligo dT 10-20 (Clontech) (0,6μg/μl), 0.8 μl 25 mM Deoxyribonucleotide triphosphate (dNTP) (Promega), 0.5 μl RNase Out (Invitrogen) and 0.5 μl Superscript III RT (200U/μl, Invitrogen) was added to a total volume of 20 μl.

RT-PCR and qPCR

Samples were incubated for two hours at 50 ºC to perform the RT-PCR reaction. Subsequently, all enzymes were destroyed at 94 ºC for 4 minutes, before cooling down to 4 ºC. Primers listed in Supplementary table 3 were used. All primers were diluted in water to a final concentration of 10 µM. Per sample, a mix was created with 12.5 µl RNase-free water, 4 µl 5x Phire buffer (Invitrogen), 0.4 µl 10 mM Deoxyribonucleotide triphosphate (dNTP) and 0.1 µl Phire III DNA polymerase (Invitrogen) to a total volume of 17 µl. Per sample 1 µl forward primer and 1 µl reverse primer was used, followed by adding 1 µl cDNA or 1 µl water as negative control to total volume of 20 µl. Reaction conditions in the PCR reaction were an initial minute at 98 ºC, then 30 cycles of 15 seconds 95 ºC, 15 seconds at 60 ºC and 15 seconds at 72 ºC, followed by 30 seconds at 72 ºC, before cooling down to 4 ºC. PCR products were analyzed on a 2% agarose (Roche) in TBE gel with 1 µl/ml ethidium bromide (Sigma). To each sample 2 µl loading reagent was added and after mixing 10 µl was put on gel, together with 5 µl marker (100bp, Invitrogen). The gel was run at constant 75 Volt for approximately 1 hour, before analyzing in UV light.

RNA sequencing

Total RNA of eight mAstro samples and 15 hAstro samples (**Supplementary M&M Table 1**) was isolated as described. RNA concentration and 260/280 ratios were measured with a NanoDrop spectrophotometer (NanoDrop 2000 Thermo Scientific). The samples were subsequently measured on a Agilent 2100 Bio Analyzer to determine RNA Integrity Number (RIN) scores. All samples had RIN scores higher than 9, so none were excluded for library construction. Illumina® TruSeq Stranded mRNA kit was used according to manufacturer's instructions in order to prepare the library, using 150 to 200ng RNA. The 3' ends were adenylylated prior to ligation of the adapters to the double stranded cDNA. In order to enrich the DNA fragments, 15 cycles of PCR were run. Quality of the product was measured on the Agilent D5000 Tape Station before sequencing.

For 8 mouse samples, sequenced fragments were aligned to mouse genome GRCm38 and annotated mouse genes (GENCODE vM12) using STAR v2.6.0 through RSEM v1.3.0 pipeline. Both expected count and Transcripts Per Kilobase Million (TPM) were obtained for in total of 50,123 genes, from which 45,193 genes located on autosomal chromosomes were extracted. Genes with TPM <1 in more than 50% of samples were excluded from the analysis, resulting in 15,290 genes. Differential expression analysis was then performed using R package DESeq2. Genes with Bonferroni corrected P-value <0.05 were considered as significantly differentially expressed genes (DEGs). We further performed gene-set enrichment analysis for DEGs using mouse Gene Ontology (GO) terms obtained from http://www.go2msig.org/cgi-bin/prebuilt.cgi?taxid=10090. The hypergeometric (upper tail) was performed for 13 significant DEGs with 15,290 background genes. After Bonferroni correction (for each GO term category separately, i.e. biological process, cellular component and molecular function), there were no significant gene sets left. All gene sets with at least two significant genes are listed in the Excel file.

For 14 human samples, sequenced fragments were aligned to human genome GRCh38 and annotated human genes (GENECODE v25) using STAR v2.6.0 through RSEM v1.3.0 pipeline. Both expected count and Transcripts Per Kilobase Million (TPM) were obtained for atotal of 63,299 genes, from which 35,414 genes located on autosomal chromosomes were extracted. Three types of DEG analyses were performed the R package DESeq2 with correcting for multiple covariates (lines, clones, replicates and cell culture subtype conditions). For each analysis, genes with TPM < 1 in 50% of the samples (not entire samples but the samples used for a specific DEG analysis) were filtered out. Only genes located on autosomal chromosomes were analyzed. DEG analyses were performed with and without covariate factors, which included individual, the number of clones per individual, and number of differentiation repetitions per clone.

| **Genotype** | **Donor/ patient** | **Gender** | **Age** | **Mutation** | **iPSC clone** | **Astrocyte differentiation** | **BrdU assay** | **ACM OPC maturation** | **Morphological analysis** | **qPCR markers** | **Poly(I:C) test** | **RNA seq** |
|---|---|---|---|---|---|---|---|---|---|---|---|---|
| control | Control 1 | Male | 44 days | - | 49 | Control-A | | Yes | F | F + C | F + C | F |
| | Control 2 | Male | 46 days | - | 42 | Control-B | Yes | Yes | C | F + C | F + C | F + C |
| | | | | | | Control-C | Yes | | | | | |
| | | | | | 43 | Control-D | Yes | | | | | |

| | | | | | | | | | | | | |
|---|---|---|---|---|---|---|---|---|---|---|---|---|
| | | | | | | Control-E | Yes | Yes | F + C | F + C | F + C | F + C |
| | Control 3 | Male | 74 days | - | 88 | Control-F | | Yes | F + C | F + C | F + C | F + C |
| | | | | | | Control-G | Yes | | | | | |
| | | | | | | Control-H | Yes | | | | | |
| | | | | | 89 | Control-I | Yes | | | | | |
| VWM | VWM307 | Male | 3 years | Homozygous, *EIF2B5*, 1484A>G | 38 | VWM-A | | Yes | F + C | C | F + C | F + C |
| | | | | | | VWM-B | Yes | Yes | C | C | F + C | F + C |
| | | | | | | VWM-C | Yes | | | | | |
| | | | | | 39 | VWM-D | | Yes | F + C | C | F + C | F + C |
| | | | | | | VWM-E | Yes | | | | | |
| | | | | | | VWM-F | Yes | | | | | |
| | VWM 389 | Female | 9 years | Homozygous, *EIF2B5*, 806G>A | 187 | VWM-G | Yes | | | | | |
| | | | | | 188 | VWM-H | | Yes | F + C | C | F + C | F + C |
| | | | | | | VWM-I | Yes | | | | | |
| | | | | | | VWM-J | Yes | | | | | |

**Supplementary M&M table 1. Origin and differentiation of human iPSC lines with an indication of which assays were performed on each differentiation. Yes = performed on hAstro (no subtype); F = FBS-hAstro; C = CNTF-hAstro.**

| **Name** | **Species** | **Company** | **Number** | **Dilution** |
|---|---|---|---|---|

| | | | | |
|---|---|---|---|---|
| GFAP | Mouse | Sigma | G3893 | 1:1000 |
| GFAP | Rabbit | DAKO | Z0334 | 1:1000 |
| Nestin | Mouse | BD biosciences | 611658 | 1:500 |
| S100β | Rabbit | ProteinTech | 15146-1-AP | 1:1000 |
| Olig2 (for mouse) | Rabbit | gift of J.H. Alberta, Harvard University, Boston, Massachusetts, USA | | 1:500 |
| Olig2 (for human) | Rabbit | Millipore | AB9610 | 1:500 |
| Sox9 | Rabbit | Cell Signalling | 82630 | 1:500 |
| MBP | Mouse | Covance | SMI-99P | 1:2000 |
| CD44 | Mouse | Hybridomabank | H4C4 | 1:250 |
| Id3 | Rabbit | Cell Signalling | 9837 | 1:250 |
| Ezrin | Mouse | Santa Cruz | sc-32759 | 1:450 |
| MOG | Mouse | Millipore | MAB5680 | 1:500 |
| OCT3/4 | Mouse | Santa Cruz | sc-5279 | 1:1000 |
| Nanog | Rabbit | Abcam | AB80892 | 1:1000 |
| Lin28a | Rabbit | Cell Signaling | 3978s | 1:1000 |
| β-Tubulin-III | Mouse | R&D Systems | MAB1195 | 1:1000 |
| α-smooth muscle actin | Mouse | Progen | 61001 | 1:1000 |
| α -fetoprotein | Mouse | R&D Systems | MAB1368 | 1:1000 |
| Tra-1-60 | Mouse | Santa Cruz | SC-21705 | 1:200 |
| SSEA4 | Mouse | Hybridomabank | (SSEA-4)-s | 1:50 |

| | | | | |
|---|---|---|---|---|
| BrdU | Rat | Abcam | ab6326 | 1:250 |

**Supplementary M&M table 2. Antibodies used for ICC.**

| **Gene name** | **Forward** | **Reverse** | **Assay** |
|---|---|---|---|
| *mGFAP* | GGGACAACTTTGCACAGGAC | GGCCACATCCATCTCCAC | RT-PCR/qPCR |
| *mS100β* | GGACACTGAAGCCAGAGAGG | TGGAAGTCACACTCCCCATC | RT-PCR/qPCR |
| *mGlast* | GAGCTACCTGCTGGGGAAT | TCCTTGGTGAGGCTCTGAAC | RT-PCR/qPCR |
| *mGLT1* | GTGGACTGGCTGCTGGATAG | TGGCTGAGAATCGGGTCATT | RT-PCR/qPCR |
| *mVimentin* | CGCTTTGCCAACTACATCGA | CCTCCTGCAATTTCTCTCGC | RT-PCR/qPCR |
| *mNestin* | CTGCAGGCCACTGAAAAGTT | GTGCTGGTCCTCTGGTATCC | RT-PCR/qPCR |
| *mGAPDH* | CGTCCCGTAGACAAAATGGT | CACCCCATTTGATGTTAGTGG | RT-PCR/qPCR |
| *mOlig2* | CCGAAAGGTGTGGATGCTTA | CACAGTCCCTCCTGTGAAGC | RT-PCR/qPCR |
| *mEif4G2* | ATTCTTCGTTGTCAAGCCGCCAAAGTGGAG | AGTTGTTTGCTGCGGAGTTGTCATCTCGTC | RT-PCR |
| *mNanog* | CAGGTGTTTGAGGGTAGCTC | CGGTTCATCATGGTACAGTC | RT-PCR |
| *mOCT3/4* | TCTTTCCACCAGGCCCCCGGCTC | TGCGGGCGGACATGGGGAGATCC | RT-PCR |
| *mSox2* | TAGAGCTAGACTCCGGGCGATGA | TTGCCTTAAACAAGACCACGAAA | RT-PCR |
| *mKLF4* | GCGAACTCACACAGGCGAGAAACC | TCGCTTCCTCTTCCTCCGACACA | RT-PCR |
| *mC-myc* | TGACCTAACTCGAGGAGGAGCTGGAATC | AAGTTTGAGGCAGTTAAAATTATGGCTGA AGC | RT-PCR |
| *mBrachyury* | ATG CCA AAG AAA GAA ACG AC | AGAGGCTGTAGAACATGATT | RT-PCR |
| *mMap2* | CATCGCCAGCCTCAGAACAAACAG | TGTACATTTCCGCCCCCAGCAG | RT-PCR |

| | | | |
|---|---|---|---|
| *mGata-6* | ACCTTATGGCGTAGAAATGCTGAGGGTG | CTGAATACTTGAGGTCACTGTTCTCGGG | RT-PCR |
| *hNANOG* | CAGCCCCGATTCTTCCACCAGTCCC | TGGAAGGTTCCCAGTCGGGTTCACC | RT-PCR |
| *hOCT3/4* | GACAGGGGGAGGGGAGGAGCTAGG | CTTCCCTCCAACCAGTTGCCCCAAAC | RT-PCR |
| *hSOX2* | GGGAAATGGGAGGGGTGCAAAAGAGG | TTGCGTGAGTGTGGATGGGATTGGTG | RT-PCR |
| *hC-MYC* | GCGTCCTGGGAAGGGAGATCCGGAGC | TTGAGGGGCATCGTCGCGGGAGGCTG | RT-PCR |
| *hTDGF* | TGCTGCTCACAGGGCCCGATACTTC | TCCTTTCGAGCTCAGTGCACCACAAAAC | RT-PCR |
| *hUTF* | CAGATCCTAAACAGCTCGCAGAAT | GCGTACGCAAATTAAAGTCCAGA | RT-PCR |
| *hDNMT3B* | CAGGAGACCTACCCTCCACA | TGTCTGAATTCCCGTTCTCC | RT-PCR |
| *hREX1* | GCTGACCACCAGCACACTAGGC | TTTCTGGTGTCTTGTCTTTGCCCG | RT-PCR |
| *hSALL4* | GCCGTGAAGACCAATGAGAT | CTCCTTCCACGCAAGTTCTC | RT-PCR |
| *hDPPA4* | GGAGCCGCCTGCCCTGGAAAATTC | TTTTTCCTGATATTCTATTCCCAT | RT-PCR |
| *hDPPA2* | CCGTCCCCGCAATCTCCTTCCATC | ATGATGCCAACATGGCTCCCGGTG | RT-PCR |
| *hESG1* | ATATCCCGCCGTGGGTGAAAGTTC | ACTCAGCCATGGACTGGAGCATCC | RT-PCR |
| *hMAP2* | CAGGTGGCGGACGTGTGAAAATTGAGAGTG | CACGCTGGATCTGCCTGGGGACTGTG | RT-PCR |
| *hSOX17* | GTGTGAATCTCCCCGACAG | TGTAACACTGCTTCTGGCC | RT-PCR |
| *hCD31* | AACAGTGTTGACATGAAGAGCC | TGTAAAACAGCACGTCATCCTT | RT-PCR |
| *hCD34* | CCTAAGTGACATCAAGGCAGAA | GCAAGGAGCAGGGAGCATA | RT-PCR |
| *hBRACHYURY 7* | GCCCTCTCCCTCCCCTCCACGCACAG | CGGCGCCGTTGCTCACAGACCACAGG | RT-PCR |
| *hSox9* | CAAGCTCTGGAGACTTCTGAACG | CCCGTTCTTCACCGACTTCCTC | RT-PCR/qPCR |
| *hALDH1L1* | TGCAGCCAATTCATCCCCAT | CTCCGTGAATGAGGGTCCAG | RT-PCR/qPCR |
| *hSLC1A2* | GCATCTACGGAAGGTGCCAA | TTGGGTTCCTCTGAGCCAAG | RT-PCR/qPCR |

| | | | |
|---|---|---|---|
| *hMLC1* | CCTAGTGGCCTGCTTTCCAA | GGACCTCCACCAGACACTTG | RT-PCR/qPCR |
| *hSLC25A18* | GGTGCCACTCTCCTCAGAGAC | TTGAACCCCAGGTTGTTAAGGT | RT-PCR/qPCR |
| *heIF4G2 (Control)* | ATTCTTCGTTGTCAAGCCGCCAAAGTGGAG | AGTTGTTTGCTGCGGAGTTGTCATCTCGTC | RT-PCR |
| *hSDHA (Control)* | CCAGGGAAGACTACAAGGTGCGGA | AGGGTGTGCTTCCTCCAGTGCT | RT-PCR/qPCR |
| *hNESTIN* | CAGGAGAAACAGGGCCTACA | TAAGAAAGGCTGGCACAGGT | RT-PCR/qPCR |
| *hBLBP* | GGATTGGGAGGAACTCGACC | CCCACGCCTAGAGCCTTCAT | RT-PCR/qPCR |
| *hS100β* | GGTGAGACAAGGAAGAGGATGT | ACAGGAAAGGTTTGGCTGCT | RT-PCR/qPCR |
| *hALDOC* | CCATGCCTGTCCCATCAAGT | TGCAAGCCCATTCACCTCAG | RT-PCR/qPCR |
| *hAQP4* | AAGGCGGTGGGGTAAGTGTG | CACTGGGCTGCGATGTAGAA | RT-PCR/qPCR |
| *hGLAST* | ACATGAAGGAACAGGGGCAG | ACCCAAGGGTTTTTCCGTGT | RT-PCR |
| *hGFAP* | GCAGATTCGAGAAACCAGCC | GAGGGCGATGTAGTAGGTGC | RT-PCR/qPCR |
| *hCD44* | TTACAGCCTCAGCAGAGCAC | AGGTGGAGCTGAAGCATTGA | RT-PCR/qPCR |
| *hEiF4G2* | AGGACCGCATGTTGGAGATT | TGAGGGGATGGATCCAACTTT | qPCR |
| *hIL6* | ACCCCCAGGAGAAGATTCCA | GATGCCGTCGAGGATGTACC | qPCR |
| *hLCN2* | TCACCCTCTACGGGAGAACC | GGGACAGGGAAGACGATGTG | qPCR |
| *hFKPB5* | AGGCTGCCATTGTCAAAGAGA | CATACTGAATCACCGCCTGC | qPCR |
| *hFLBN5* | TGCCAGGAATAAAAAGGATACTCAC | ACTGGCGATCCAGGTCAAAG | qPCR |
| *hPTX3* | GACTCCATCCCACTGAGGAC | CAGCATGCGCTCTCTCATCT | qPCR |
| *hTLR3* | AGAAAGGGACTTTGAGGCGG | TGTTGAACTGCATGATGTACCTTG | qPCR |
| *hSerping1* | TCTCCTAACACTACCCCGCA | CAGCCCACACAGGTTAAGGT | qPCR |

**Supplementary M&M table 3. Primers used in PCR analyses.**

#### Supplementary references


Dooves S, Bugiani M, Postma NL, Polder E, Land N, Horan ST*, et al.* Astrocytes are central in the pathomechanisms of vanishing white matter. J Clin Invest 2016; 126(4): 1512-24.

Nadadhur AG, Leferink PS, Holmes D, Hinz L, Cornelissen-Steijger P, Gasparotto L*, et al.* Patterning factors during neural progenitor induction determine regional identity and differentiation potential in vitro. Stem Cell Res 2018; 32: 25-34.

Takahashi K, Yamanaka S. Induction of pluripotent stem cells from mouse embryonic and adult fibroblast cultures by defined factors. Cell 2006; 126(4): 663-76.

Warlich E, Kuehle J, Cantz T, Brugman MH, Maetzig T, Galla M*, et al.* Lentiviral vector design and imaging approaches to visualize the early stages of cellular reprogramming. Mol Ther 2011; 19(4): 782-9.
