## Supplementary figures and images for "Human and mouse iPSC-derived astrocyte subtypes reveal vulnerability in Vanishing White Matter"

### Supplemental Figure 1

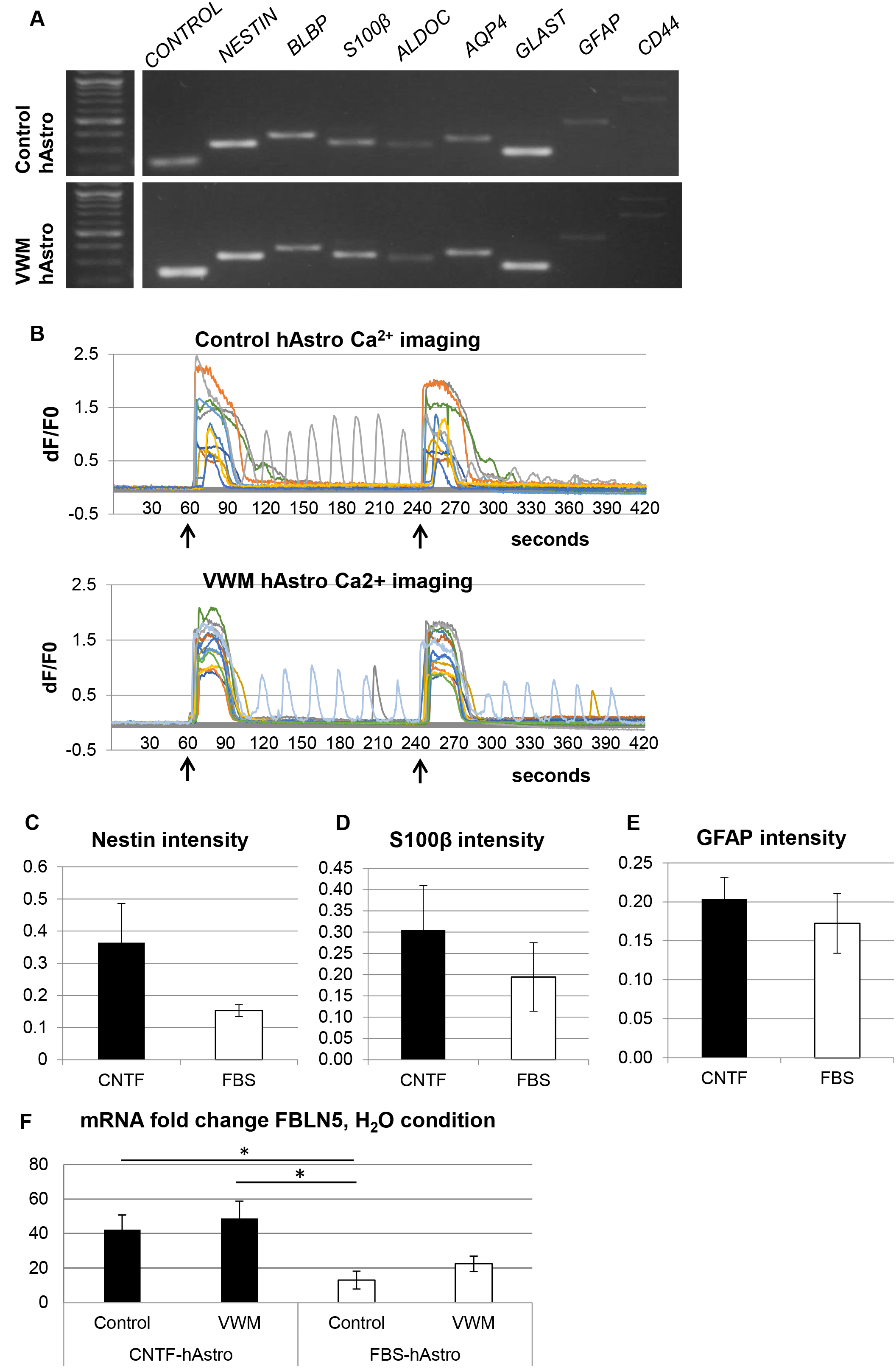
